# Multi-organ single-cell RNA-sequencing reveals early hyperglycaemia responses that converge on fibroblast dysregulation

**DOI:** 10.1101/2023.08.07.552307

**Authors:** Adam T. Braithwaite, Naveed Akbar, Daniela Pezzolla, Daan Paget, Thomas Krausgruber, Christoph Bock, Ricardo Carnicer, Robin P. Choudhury

## Abstract

Diabetes causes a range of complications that can affect multiple organs. Hyperglycaemia is an important driver of diabetes-associated complications, mediated by biological processes such as dysfunction of endothelial cells, fibrosis and alterations in leukocyte number and function. Here, we dissected the transcriptional response of key cell types to hyperglycaemia across multiple tissues using single-cell RNA-seq (scRNA-seq) and identified conserved, as well as organ-specific, changes associated with diabetes complications. By studying an early timepoint of diabetes, we focus on biological processes involved in the initiation of the disease, before the later organ-specific manifestations had supervened. We used a mouse model of type 1 diabetes and performed scRNA-seq on cells isolated from the heart, kidney, liver and spleen of streptozotocin-treated and control mice after 8 weeks and assessed differences in cell abundance, gene expression, pathway activation and cell signalling across organs and within organs. In response to hyperglycaemia, endothelial cells, macrophages and monocytes displayed organ-specific transcriptional responses, whereas fibroblasts showed similar responses across organs, exhibiting a myofibroblast-like phenotype with altered metabolic gene expression and increased differentiation of myeloid-derived fibroblasts. Further, we found evidence of endothelial dysfunction in the kidney, and of endothelial to mesenchymal transition in streptozotocin-treated mouse organs. In summary, our study represents the first single-cell and multi-organ analysis of early dysfunction in type 1 diabetes-associated hyperglycaemia, and our large-scale dataset (comprising 67,611 cells) will serve as a starting point, reference atlas, and resource for further investigating the events leading to early diabetic disease.

## Introduction

Hyperglycaemia is central to the diagnosis, monitoring and treatment in patients with both type 1 and type 2 diabetes. Exposure to blood glucose elevation is shared by all organs and tissues, but the loss of function leading to clinical manifestations are, naturally, organ specific. However, some underlying molecular and cellular pathologies could be shared. For instance, high glucose levels can exacerbate disease (1) by initiating the nonenzymatic glycation of proteins and lipoproteins, leading to activation of the receptor for advanced glycation end products (RAGE) (Stirban et al., 2013), (2) by driving oxidative stress (Giacco and Brownlee, 2010) and (3) by promoting inflammation (Tsalamandris et al., 2019). Several cell types contribute to the progression of diabetes-associated complications and include dysfunction of endothelial cells (ECs) (Abebe and Mozaffari, 2010), fibrosis / deposition of extracellular matrix (ECM) by fibroblasts, (Ban and Twigg, 2008) and alterations in generation of leukocytes, and their programming and recruitment (Bajpai and Tilley, 2018; Edgar et al., 2021; Szabo et al., 2007).

Alterations in cellular differentiation and functional states are observed in many different diseases and contribute to diabetes-associated complications. For example, activated, pro-fibrotic cells can be derived from other transitioning cell types via a biological process called epithelial to mesenchymal transition (EMT), where a polarised epithelial cell, which normally interacts with basement membrane via its basal surface, assumes a mesenchymal cell phenotype, characterised by enhanced migratory capacity, invasiveness, resistance to apoptosis, and greatly increased production of ECM components. Similarly, circulating bone marrow-derived myeloid cells, such as monocytes or macrophages, have been shown to transition into fibrocytes or fibroblasts that contribute to the development of fibrosis-related complications of diabetes (Abu El-Asrar et al., 2015; Yan et al., 2016). Furthermore, in endothelial to mesenchymal transition (EndoMT), an EC undergoes a series of molecular events that changes its phenotype toward a mesenchymal cell with myofibroblast or smooth muscle cell function. EndoMT causes loss of tight junctions in glomeruli leading to increased endothelial permeability and albuminuria (Loeffler and Wolf, 2015; Wu et al., 2018).

Given the potential for shared early cellular pathologies secondary to hyperglycaemia, and the importance of ECs, fibroblasts, macrophages and monocytes across organs affected by these pathologies, we hypothesised that processes of disease initiation might be shared between organs, with subsequent divergence and clinical manifestation according to organ-specific composition and function. Accordingly, we performed a comprehensive characterisation of cell populations at an early time point prior to overt pathology, and determined single-cell RNA-seq (scRNA-seq) profiles in four organs (heart, kidney, liver and spleen) associated with complications in diabetes using the streptozotocin (STZ) model of hyperglycaemia in mice. scRNA-seq is a powerful approach for determining cellular composition and transcriptional phenotypes, allowing comparisons between source conditions such as developmental stage; organ of origin and pathological state (Hwang et al., 2018). While several studies have utilised this technology to characterise cells from affected organs individually; particularly the kidney (Wilson et al., 2022, 2019), none, to our knowledge have attempted an integrated systematic analysis of hyperglycaemia-related changes across disease-relevant organs.

Our study uncovered changes in cellular developmental and functional programming with plausible roles in the initiation of the complications of diabetes. Overall, we identified organ-specific as well as shared pathological mechanisms, providing potential targets for preventative and therapeutic interventions for diabetes-associated complications.

## Results

### Multi-organ single cell transcriptome profiles from hyperglycaemic and control mice

To study cell composition and gene regulation in diabetes, we used the STZ mouse model of hyperglycaemia and specifically focused on an early timepoint, to investigate and characterise the initiation of diabetes-related pathology. STZ administration for five consecutive days led to the production of non-fasted blood glucose levels (>20 mmol/L) and increased terminal glucose at the 8-week timepoint (mean 26.8 ± 6.8 in STZ mice (n=4) *vs* 7.9 ± 0.4 mmol/L in control mice (n=3), *p* = 0.005; data not shown) associated with diabetic disease. We also observed an increase in organ masses upon STZ treatment in the liver (liver/body mass ratio: mean 55.4 *vs* 42.8 mg/g, *p* = 0.027) and kidney (kidney/body mass ratio: mean 13.59 *vs* 11.15 mg/g, *p* = 0.047; data not shown).

To unravel the gene regulatory programs across organs, we isolated cells from the heart ventricles, kidney, liver and spleen of control and STZ-treated mice and performed scRNA-seq. Cell quality was assessed by taking percentage of mitochondria, number of genes detected per cell, and number of Unique Molecular Identifiers (UMI) into account (Figure S1A–C). Cells were assigned to an individual tissue according to their hashtag oligo (HTO) read counts and doublets (cells with multiple HTO barcodes) discarded during this processing step (Figure S1D–G). We integrated the data into a joint map and performed clustering according to HTO barcode levels to confirm distinct cluster formation of singlet cells, with doublet cells located between them (Figure S1H). From each mouse a total of 7,902 to 11,651 singlet cells were included in the subsequent analyses, resulting in a total of 40,710 cells from STZ treated mice and 26,901 cells from control mice.

### Cell type profiles are organ-specific but relatively stable across conditions

Single-cell transcriptome profiles from all organs of the two experimental groups were integrated, and cells were clustered by gene expression (Figure 1A). Cell clusters were annotated based on organ specificity and differential expression of key marker genes (vs. other clusters) that were conserved across STZ and control groups (Figure 1B). The marker genes reflected well-known cell surface proteins in addition to mRNA / single cell-specific markers extracted from published annotations (summarised in Table 1; further details available in Supplementary Data 1). The identified cell types and cell subsets were present in the STZ and control group (Figure 1A) indicating that the treatment had no overt cell-type specific effect. We explored the relationships between clusters by building a classification hierarchy in which transcriptionally similar clusters assumed adjacencies on a dendrogram (Figure 1C). As expected, this analysis showed distinct branches for myeloid, lymphoid and mesenchymal cell types.

**Figure 1.**
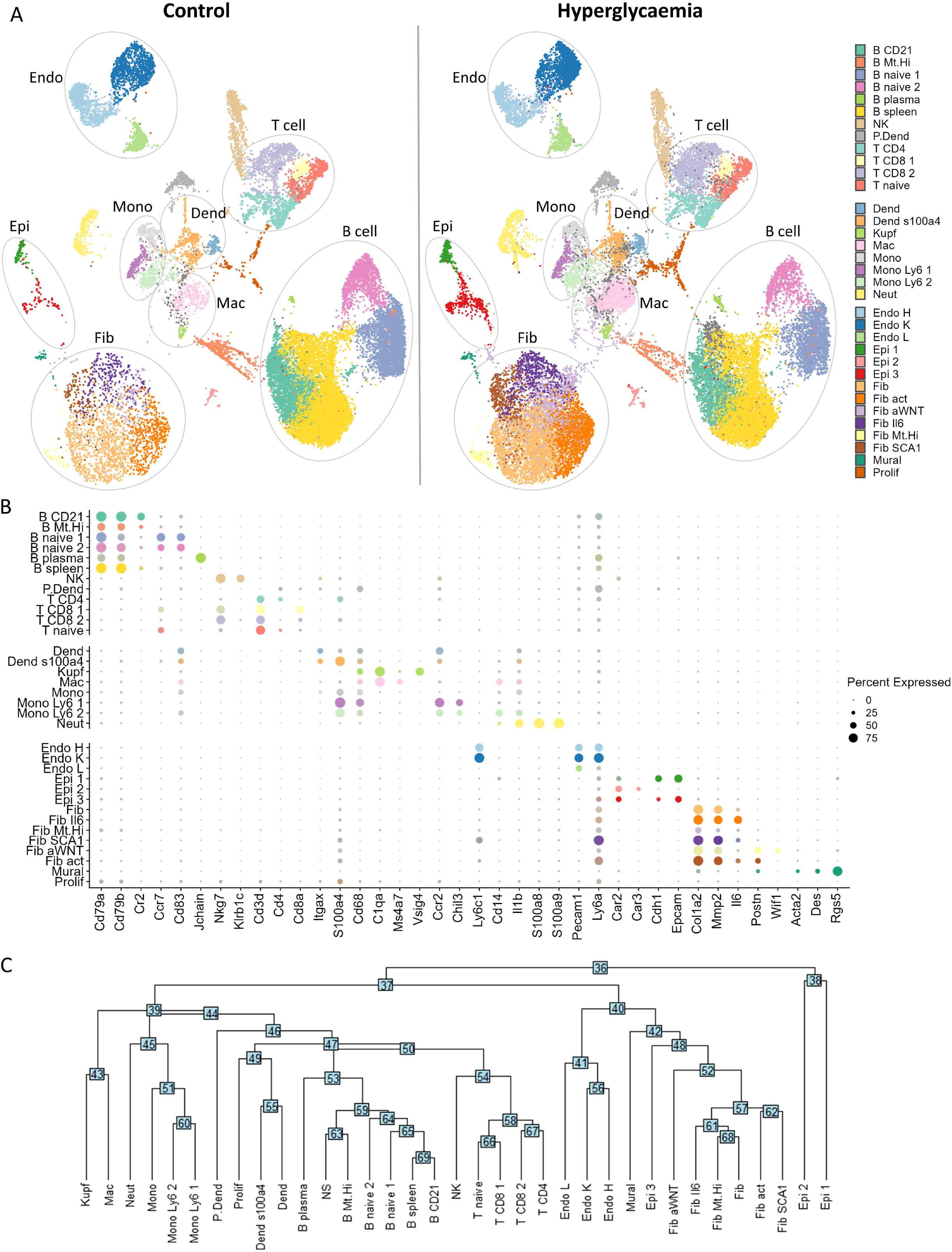
Multi-organ single cell transcriptome profiles from hyperglycaemic and control mice. (A) Single cell profiles from all organs were integrated and cells clustered. t-distributed stochastic neighbour embedding (tSNE) plots show a matched number of cells from each condition. (B) Average expression of key marker genes that were used to annotate specific cell clusters and sub-clusters. (C) The relative functional proximity of clusters was demonstrated by building a classification hierarchy with transcriptionally similar clusters adjacent on a tree. Dend, myeloid dendritic cell; Endo, endothelial cell; Epi, epithelial cell; Fib, fibroblast; Mac, macrophage; Mono, monocyte.

**Table 1:**
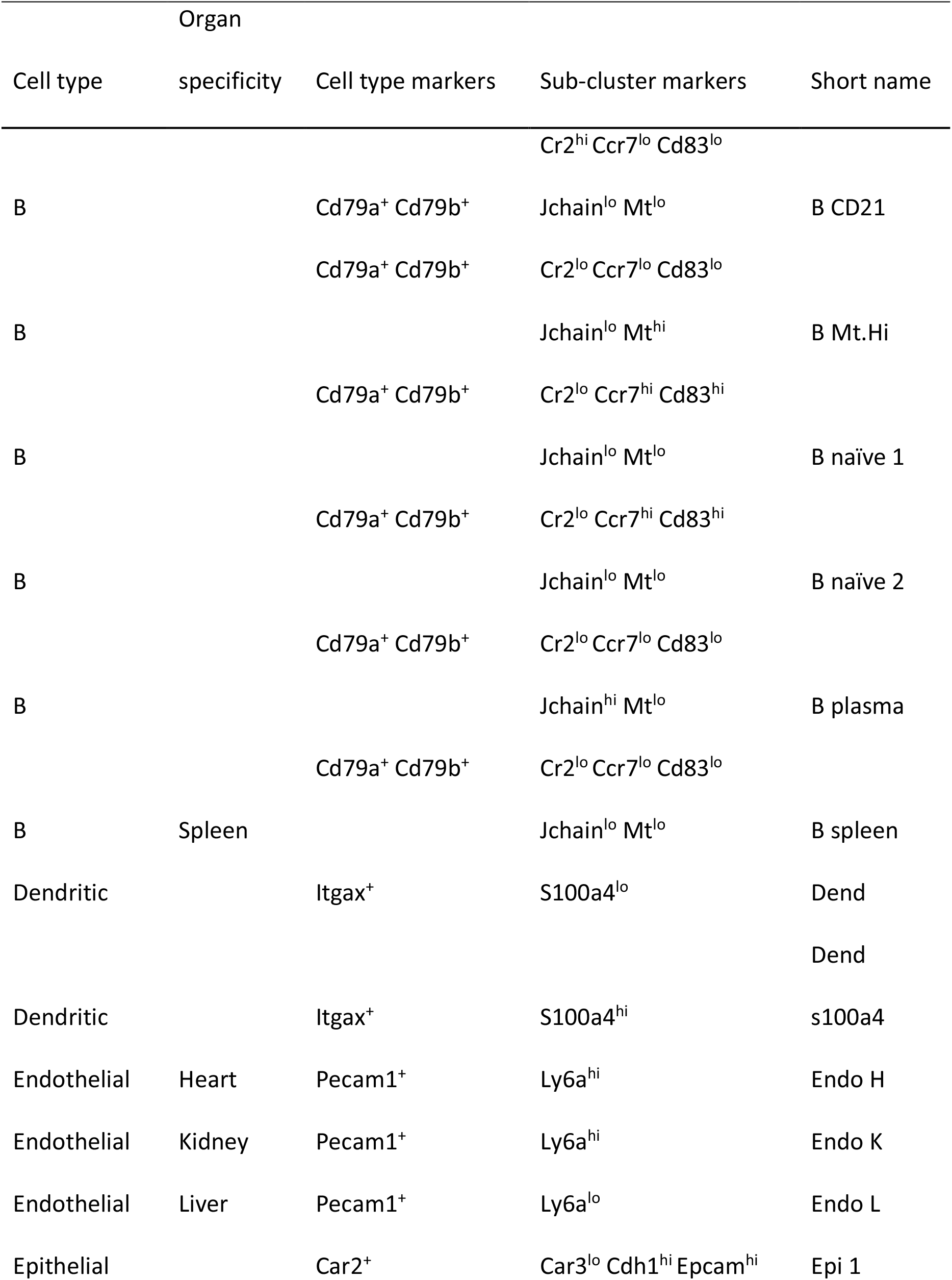
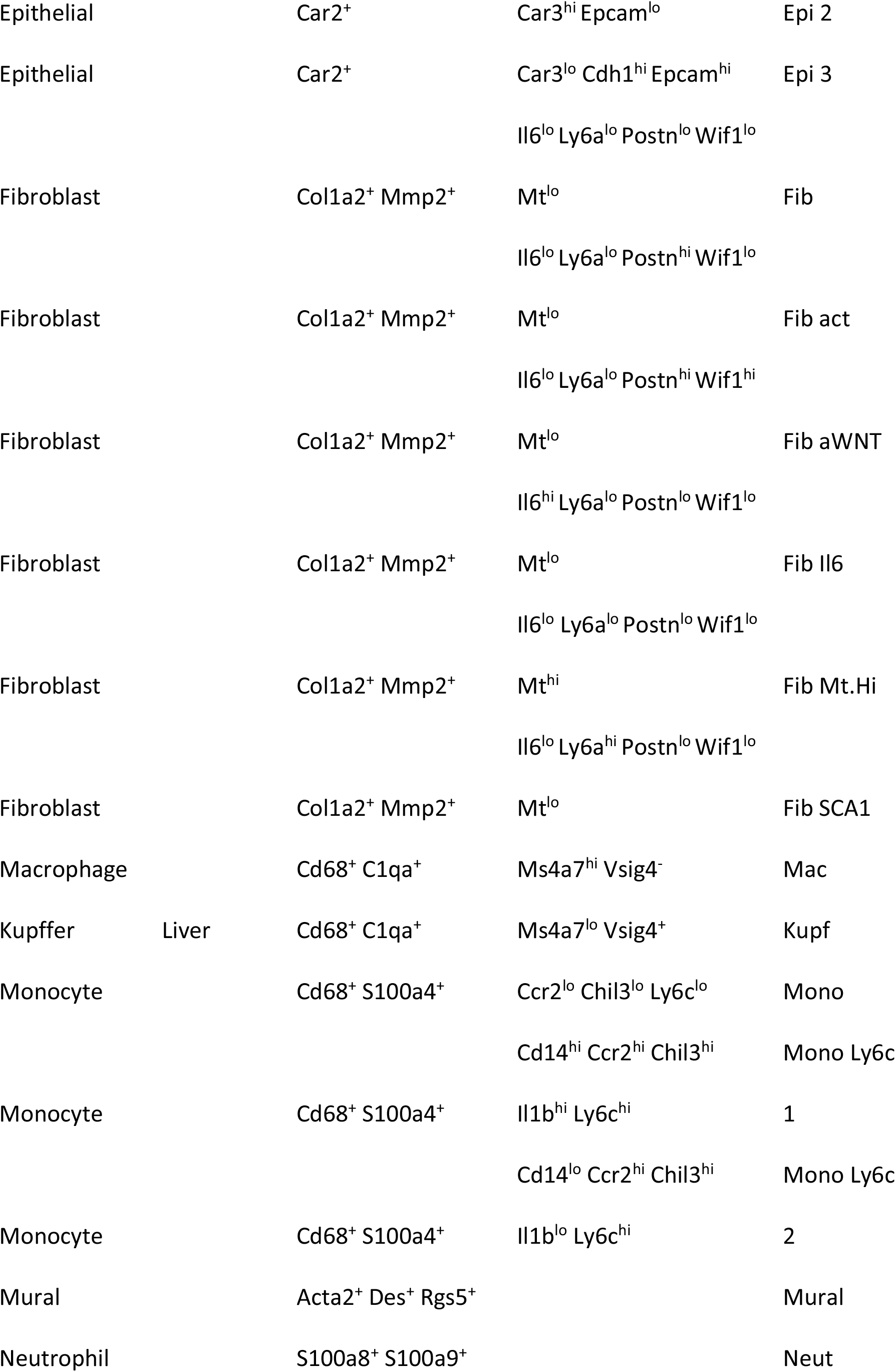
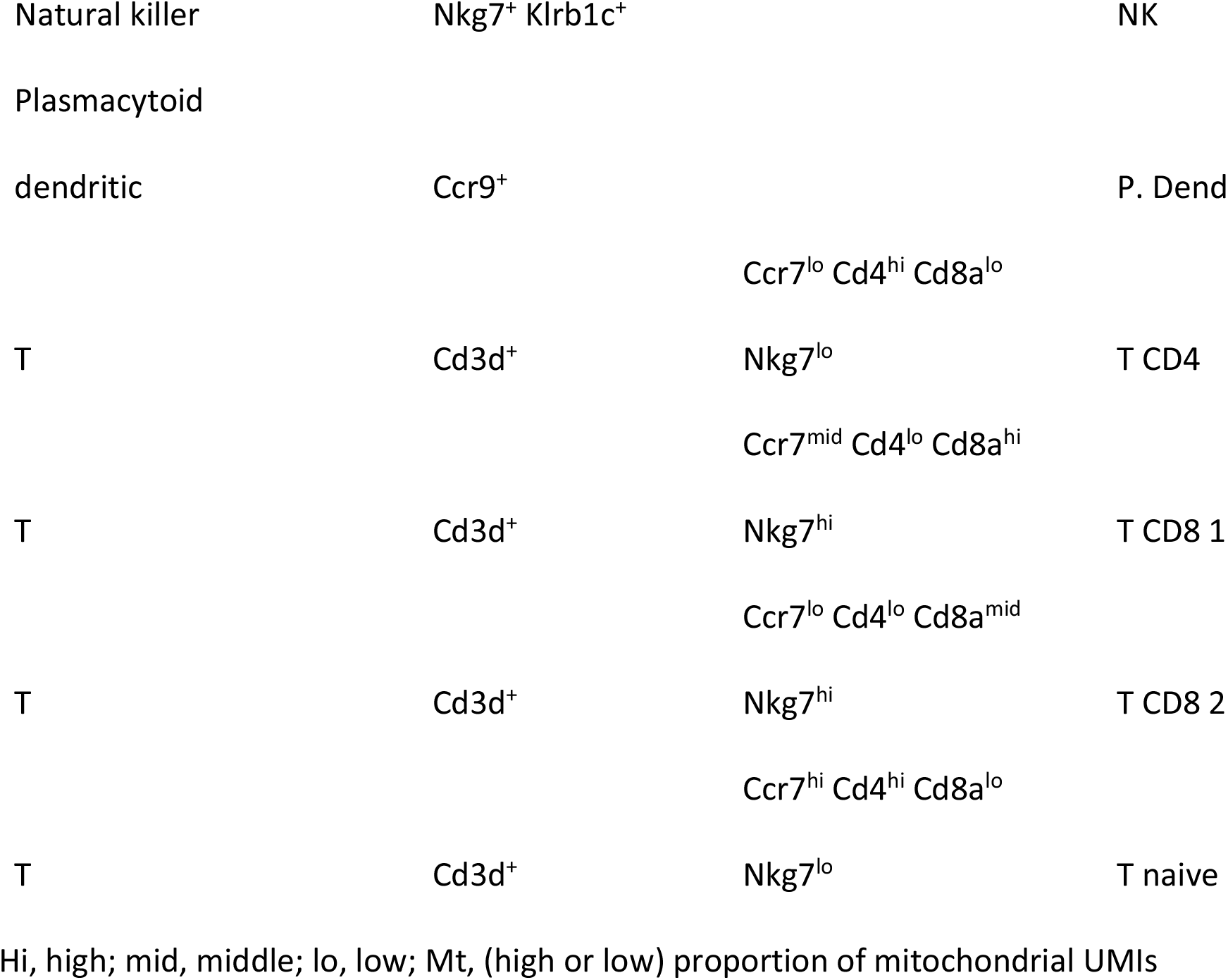
Annotation of cell types and sub-clusters with specific marker genes.

| Organ |  |  |  |  |
| --- | --- | --- | --- | --- |
| Cell type | specificity | Cell type markers | Sub-cluster markers | Short name |
|  |  |  | Cr2<sup>hi</sup> Ccr7<sup>lo</sup> Cd83<sup>lo</sup> |  |
| B |  | Cd79a<sup>+</sup> Cd79b<sup>+</sup> | Jchain<sup>lo</sup> Mt<sup>lo</sup> | B CD21 |
|  |  | Cd79a<sup>+</sup> Cd79b<sup>+</sup> | Cr2<sup>lo</sup> Ccr7<sup>lo</sup> Cd83<sup>lo</sup> |  |
| B |  |  | Jchain<sup>lo</sup> Mt<sup>hi</sup> | B Mt.Hi |
|  |  | Cd79a<sup>+</sup> Cd79b<sup>+</sup> | Cr2<sup>lo</sup> Ccr7<sup>hi</sup> Cd83<sup>hi</sup> |  |
| B |  |  | Jchain<sup>lo</sup> Mt<sup>lo</sup> | B naïve 1 |
|  |  | Cd79a<sup>+</sup> Cd79b<sup>+</sup> | Cr2<sup>lo</sup> Ccr7<sup>hi</sup> Cd83<sup>hi</sup> |  |
| B |  |  | Jchain<sup>lo</sup> Mt<sup>lo</sup> | B naïve 2 |
|  |  | Cd79a<sup>+</sup> Cd79b<sup>+</sup> | Cr2<sup>lo</sup> Ccr7<sup>lo</sup> Cd83<sup>lo</sup> |  |
| B |  |  | Jchain<sup>hi</sup> Mt<sup>lo</sup> | B plasma |
|  |  | Cd79a<sup>+</sup> Cd79b<sup>+</sup> | Cr2<sup>lo</sup> Ccr7<sup>lo</sup> Cd83<sup>lo</sup> |  |
| B | Spleen |  | Jchain<sup>lo</sup> Mt<sup>lo</sup> | B spleen |
| Dendritic |  | Itgax<sup>+</sup> | S100a4<sup>lo</sup> | Dend |
|  |  |  |  | Dend |
| Dendritic |  | Itgax<sup>+</sup> | S100a4<sup>hi</sup> | s100a4 |
| Endothelial | Heart | Pecam1<sup>+</sup> | Ly6a<sup>hi</sup> | Endo H |
| Endothelial | Kidney | Pecam1<sup>+</sup> | Ly6a<sup>hi</sup> | Endo K |
| Endothelial | Liver | Pecam1<sup>+</sup> | Ly6a<sup>lo</sup> | Endo L |
| Epithelial |  | Car2<sup>+</sup> | Car3<sup>lo</sup> Cdh1<sup>hi</sup> Epcam<sup>hi</sup> | Epi 1 |
| Epithelial |  | Car2<sup>+</sup> | Car3<sup>hi</sup> Epcam<sup>lo</sup> | Epi 2 |
| Epithelial |  | Car2<sup>+</sup> | Car3<sup>lo</sup> Cdh1<sup>hi</sup> Epcam<sup>hi</sup> | Epi 3 |
|  |  |  | Il6<sup>lo</sup> Ly6a<sup>lo</sup> Postn<sup>lo</sup> Wif1<sup>lo</sup> |  |
| Fibroblast |  | Col1a2<sup>+</sup> Mmp2<sup>+</sup> | Mt<sup>lo</sup> | Fib |
|  |  |  | Il6<sup>lo</sup> Ly6a<sup>lo</sup> Postn<sup>hi</sup> Wif1<sup>lo</sup> |  |
| Fibroblast |  | Col1a2<sup>+</sup> Mmp2<sup>+</sup> | Mt<sup>lo</sup> | Fib act |
|  |  |  | Il6<sup>lo</sup> Ly6a<sup>lo</sup> Postn<sup>hi</sup> Wif1<sup>hi</sup> |  |
| Fibroblast |  | Col1a2<sup>+</sup> Mmp2<sup>+</sup> | Mt<sup>lo</sup> | Fib aWNT |
|  |  |  | Il6<sup>hi</sup> Ly6a<sup>lo</sup> Postn<sup>lo</sup> Wif1<sup>lo</sup> |  |
| Fibroblast |  | Col1a2<sup>+</sup> Mmp2<sup>+</sup> | Mt<sup>lo</sup> | Fib Il6 |
|  |  |  | Il6<sup>lo</sup> Ly6a<sup>lo</sup> Postn<sup>lo</sup> Wif1<sup>lo</sup> |  |
| Fibroblast |  | Col1a2<sup>+</sup> Mmp2<sup>+</sup> | Mt<sup>hi</sup> | Fib Mt.Hi |
|  |  |  | Il6<sup>lo</sup> Ly6a<sup>hi</sup> Postn<sup>lo</sup> Wif1<sup>lo</sup> |  |
| Fibroblast |  | Col1a2<sup>+</sup> Mmp2<sup>+</sup> | Mt<sup>lo</sup> | Fib SCA1 |
| Macrophage |  | Cd68<sup>+</sup> C1qa<sup>+</sup> | Ms4a7<sup>hi</sup> Vsig4<sup>-</sup> | Mac |
| Kupffer | Liver | Cd68<sup>+</sup> C1qa<sup>+</sup> | Ms4a7<sup>lo</sup> Vsig4<sup>+</sup> | Kupf |
| Monocyte |  | Cd68<sup>+</sup> S100a4<sup>+</sup> | Ccr2<sup>lo</sup> Chil3<sup>lo</sup> Ly6c<sup>lo</sup> | Mono |
|  |  |  | Cd14<sup>hi</sup> Ccr2<sup>hi</sup> Chil3<sup>hi</sup> | Mono Ly6c |
| Monocyte |  | Cd68<sup>+</sup> S100a4<sup>+</sup> | Il1b<sup>hi</sup> Ly6c<sup>hi</sup> | 1 |
|  |  |  | Cd14<sup>lo</sup> Ccr2<sup>hi</sup> Chil3<sup>hi</sup> | Mono Ly6c |
| Monocyte |  | Cd68<sup>+</sup> S100a4<sup>+</sup> | Il1b<sup>lo</sup> Ly6c<sup>hi</sup> | 2 |
| Mural |  | Acta2<sup>+</sup> Des<sup>+</sup> Rgs5<sup>+</sup> |  | Mural |
| Neutrophil |  | S100a8<sup>+</sup> S100a9<sup>+</sup> |  | Neut |

|  |  |  |  |
| --- | --- | --- | --- |
| Natural killer | Nkg7<sup>+</sup> Klrb1c<sup>+</sup> |  | NK |
| Plasmacytoid |  |  |  |
| dendritic | Ccr9<sup>+</sup> |  | P. Dend |
|  |  | Ccr7<sup>lo</sup> Cd4<sup>hi</sup> Cd8a<sup>lo</sup> |  |
| T | Cd3d<sup>+</sup> | Nkg7<sup>lo</sup> | T CD4 |
|  |  | Ccr7<sup>mid</sup> Cd4<sup>lo</sup> Cd8a<sup>hi</sup> |  |
| T | Cd3d<sup>+</sup> | Nkg7<sup>hi</sup> | T CD8 1 |
|  |  | Ccr7<sup>lo</sup> Cd4<sup>lo</sup> Cd8a<sup>mid</sup> |  |
| T | Cd3d<sup>+</sup> | Nkg7<sup>hi</sup> | T CD8 2 |
|  |  | Ccr7<sup>hi</sup> Cd4<sup>hi</sup> Cd8a<sup>lo</sup> |  |
| T | Cd3d<sup>+</sup> | Nkg7<sup>lo</sup> | T naive |

Hi, high; mid, middle; lo, low; Mt, (high or low) proportion of mitochondrial UMIs

Following integration and annotation, we investigated associations of scRNA-seq profiles with organ of origin and/or hyperglycaemia. Cells from all clusters and sub-clusters were represented in the STZ and control conditions for each organ (Figure 2A). Cell clusters were summarised by general categories (lymphoid, myeloid or other cells) to compare their relative distributions between individual mice, organs and disease groups (Figure 2B).

**Figure 2.**
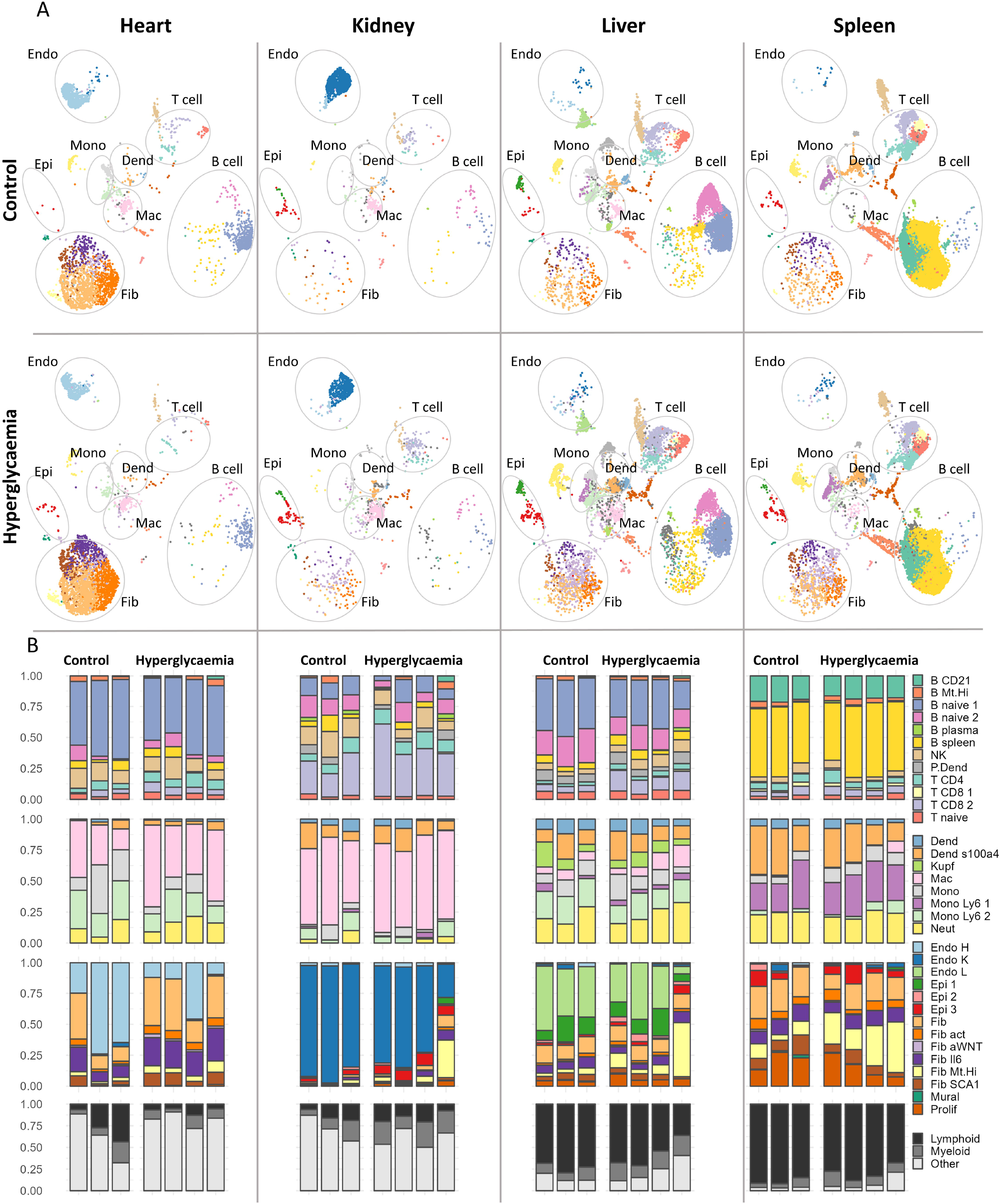
Cell type profiles are organ-specific but relatively stable across conditions. (A) Annotated cells split by tissue of origin and group. t-distributed stochastic neighbour embedding (tSNE) plots show a matched number of cells from each condition per tissue. (B) Cell sub-clusters summarised by general type (lymphoid, myeloid or other cells) and relative cell proportions are indicated in one column per mouse. Dend, myeloid dendritic cell; Endo, endothelial cell; Epi, epithelial cell; Fib, fibroblast; Mac, macrophage; Mono, monocyte.

To study the effect of glucose on cell-type-specific and organ-specific gene regulation, we composed pseudo bulk RNA-seq profiles by summing reads per gene per mouse for key cell types in each organ. Variation in gene expression for each pseudo bulk profile was summarised and visualised for the first two principal components showing an organ-specific, rather than treatment-specific, clustering for ECs, fibroblasts, macrophages and monocytes (Figure 3A). Interestingly, organ-specific clusters within fibroblasts were less distinct, indicating greater phenotypic similarity between organs than in other cell types.

**Figure 3.**
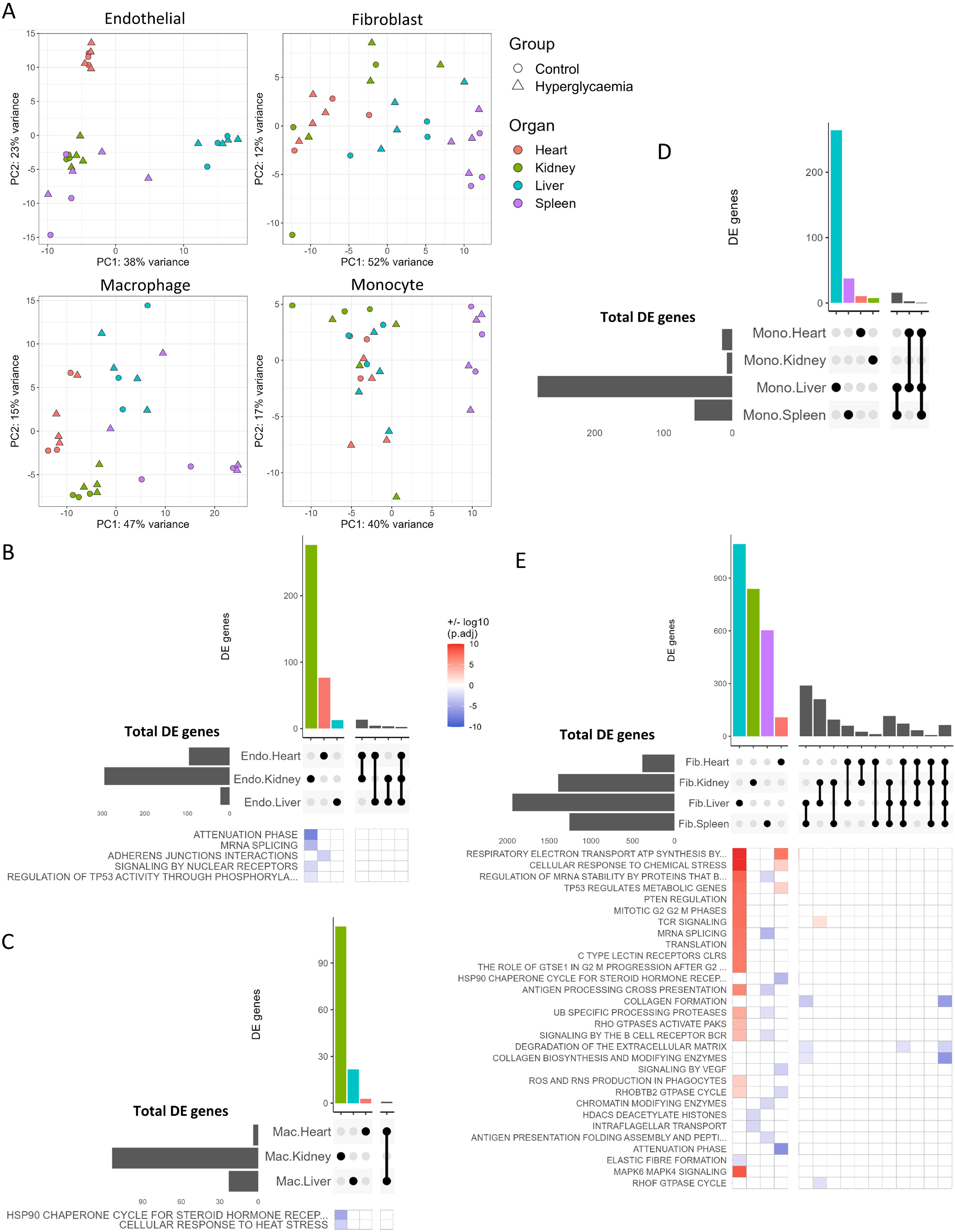
Transcriptional responses to hyperglycaemia converge on fibroblasts. (A) Pseudo bulk RNA-sequencing profiles were generated for each mouse/tissue/cell type by summing reads per gene per sample. The variation of gene expression for each pseudo bulk profile was calculated using the top 150 most variable genes and visualised for the first two principle components. Each point represents summed cells from one mouse. (B–E) For key cell types, common significant differentially expressed genes (STZ vs. control; adjusted p < 0.05 and log2FC > 0.1) were assessed across organs. Upset plots show distinct and shared differential genes from (B) endothelial cells, (C) macrophages, (D), monocytes, (E) fibroblasts, per organ. Heatmaps show +/− log10 (adjusted p-value) of significantly enriched pathways from over-representation analysis of Reactome pathways using each distinct and shared differential set of up- and downregulated genes (log2FC > 0.1, p<0.05) per organ (with no enriched pathways found in monocytes). Endo, endothelial cell; Fib, fibroblast; Mac, macrophage; Mono, monocyte.

### Transcriptional responses to hyperglycaemia converge on fibroblasts

To determine functional changes due to hyperglycaemia, we performed differential expression analysis (with p < 0.05 and FC log2 ± 0.1) of cells from STZ vs. control mice (Figure S2–5). We then summarised the number of differentially expressed genes for ECs, macrophages, monocytes and fibroblasts across organs (Figure 3B–E). In ECs, macrophages and monocytes only a minority of hyperglycaemia-responsive genes were shared across organs, indicating relative organ-specificity of cellular response to hyperglycaemia. However, in fibroblasts we detected a greater proportion of differentially expressed genes shared across organs and their transcriptome profiles changed strongly in response to hyperglycaemia. This effect was not due to greater sampling of fibroblast gene expression (due to cell abundance), as fibroblasts were not the most numerous cells in kidney, liver or spleen. Shared patterns of differential gene expression from STZ-treated mice were highest in liver and spleen fibroblasts (569 genes), followed by kidney and liver fibroblasts (455 genes). Nevertheless, a majority of differentially expressed genes in STZ mouse fibroblasts were organ-specific, which may reflect the diversity of their biological functions in different organs.

To elucidate the biological function of the transcriptional response to hyperglycaemia, we used an over-representation analysis to annotate the Reactome pathways associated with groups of differentially expressed genes that were organ-specific or shared across organs (Figure 3B–E). Several enriched pathways in STZ-treated fibroblasts were more organ-specific, including metabolism-related terms ‘TP53 regulates metabolic genes’ and ‘Respiratory electron transport, ATP synthesis by chemiosmotic coupling, and heat production by uncoupling proteins’ in STZ mouse liver and heart, but not in kidney or spleen. Related to this enrichment of metabolic pathways, we also noted an increased proportion of high-mitochondria (>10% of UMIs) fibroblasts in STZ mouse heart (Figure S6). Conversely to organ-specific responses, pathways relating to collagen/ECM (e.g. ‘Collagen formation’, ‘Degradation of the extracellular matrix’) were associated with genes downregulated in fibroblasts from all four organs. Similar pathways were also found to be downregulated in fibroblasts using a more sensitive ranked gene set enrichment analysis (Figure S2–5), confirming the robustness of the results. The observed pathway dysregulation was driven by lower expression of transcripts coding for collagen genes (including *Col1a1, Col4a4* and *Col5a1*; data not shown), however, collagen genes remained among the most highly expressed transcripts within fibroblasts of both STZ and control mice. Expression of ECM-degrading matrix metalloprotease genes (particularly *Mmp14* and *Mmp23*) were also reduced in kidney, liver and spleen fibroblasts, but not in heart fibroblasts. Suppression of matrix metalloprotease genes has been linked to an activated/myofibroblast phenotype (Howard et al., 2012). Notably the contractile marker beta actin (*Actb*) was upregulated in heart, kidney, liver and spleen fibroblasts of STZ treated mice. In contrast, the expression of classical fibroblast markers including *Pdgfra* in liver and spleen as well as *Pdgfrb* in heart, kidney, liver and spleen was decreased in the STZ treatment group.

### Hyperglycaemia induces myeloid-derived fibroblasts

In addition to shared hyperglycaemia-responsive genes, we also detected an upregulation of lysozyme 2 (*Lyz2*), a marker of fibroblasts/fibrocytes of circulating myeloid cell origin (Guerrero-Juarez et al., 2019), in fibroblasts from the STZ treatment group. This was particularly apparent in kidney and liver where *Lyz2* was the most highly upregulated gene (kidney, log2-FC 2.99, p < 0.01; liver, log2-FC 2.13, p < 0.0001). Overall, the proportion of fibroblast expressing *Lyz2* was increased in all organs upon STZ treatment compared control mice (Figure 4A), with the largest disparity in kidney (46.4% vs. 8.5% *Lyz2^+^*). To further characterise the *Lyz2^+^* fibroblasts, we classified all fibroblasts based on *Lyz2* expression and examined key fibroblast marker genes (Figure 4B). *Lyz2^+^* fibroblasts showed a reduction in *Col1a2* and *Mmp2* expression, but an increase in *Actb* expression in all organs compared to *Lyz2^−^* fibroblasts (Figure 4B). Together, these data indicate that the increased proportion of *Lyz2^+^*fibroblasts in STZ mice contributes to the apparent reduction in collagen and matrix metalloprotease gene expression (when averaged across all fibroblast populations), and at the same time drives the expression of contractile markers (e.g. beta-actin). Notably, more *Lyz2^+^* fibroblasts in kidney, liver and spleen were positive for expression of the monocyte/macrophage marker *Cd14* and macrophage marker *Cd68* than *Lyz2^−^* fibroblasts, but the expression of *Cd34* was decreased, suggesting that these cells might be macrophage-derived fibroblasts rather than monocyte-derived fibrocytes. A global differential expression analysis revealed an upregulation of complement genes (*C1qa*, *C1qb*, *C1qc*) in all four organs, MHC class II genes (*H2-Aa*, *H2-Ab1*, *H2-Eb1*) in heart, kidney and liver; and *Il1b* in heart, liver and spleen *Lyz2^+^*fibroblasts (Figure 4C). These gene expression differences were associated with differentially enriched functional pathways (Figure 4D).

**Figure 4.**
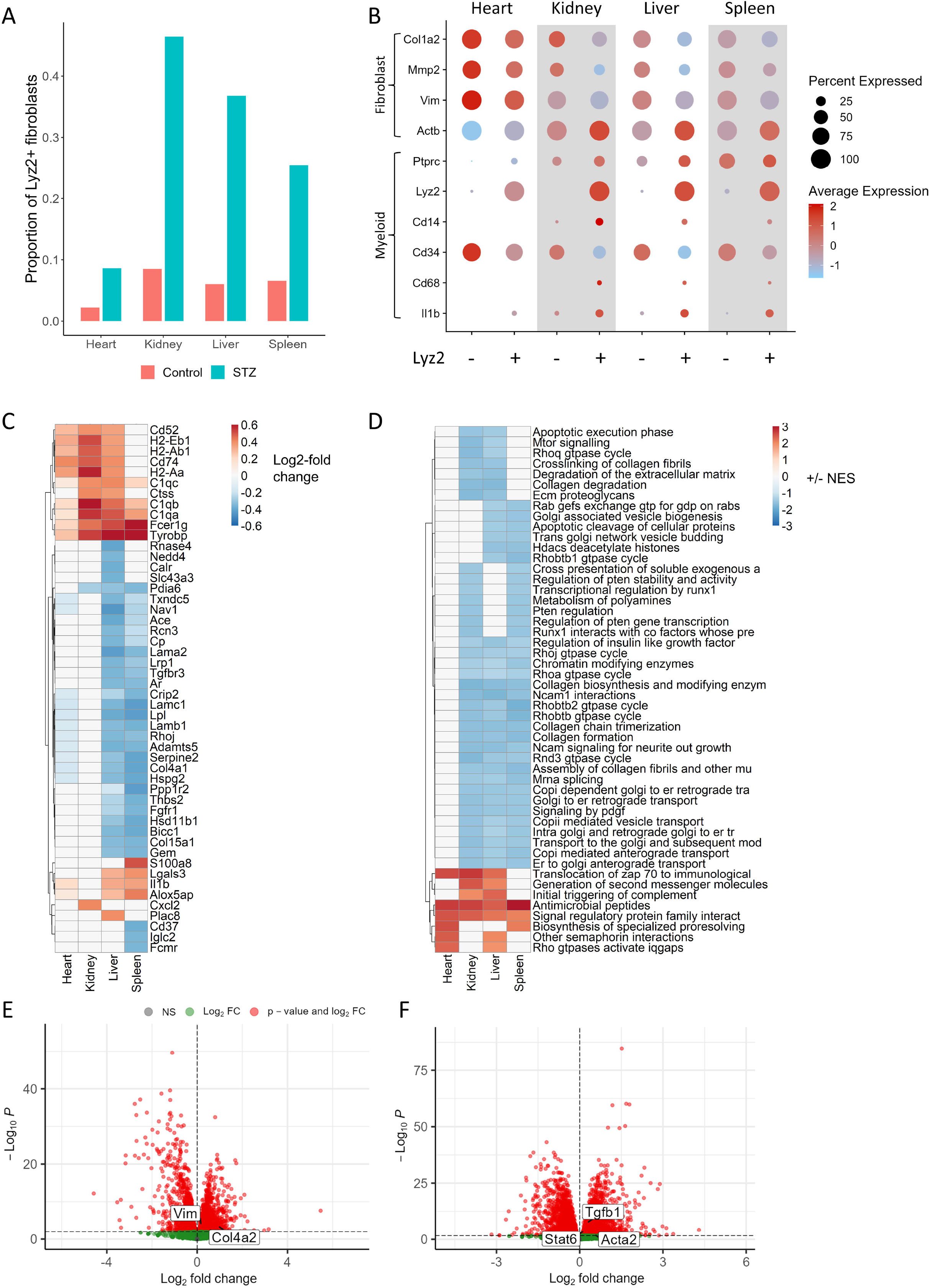
Hyperglycaemia induces myeloid-derived fibroblasts. (A) Overall proportion of Lyz2^+^ fibroblasts when combining all cells from each organ/group. (B) Dot plots representing expression of key marker genes within Lyz2^+^ and Lyz2^−^ mice fibroblasts by organ. Circle colour represents average expression across cells, and circle size represents the percentage of cells positive for gene expression. (C) Top 50 differentially expressed genes (when ranked by maximum log2FC) comparing Lyz2^+^ and Lyz2^−^ fibroblasts from each organ. Significantly differentially expressed genes (adjusted p<0.05) are coloured, with colour representing log2FC (trimmed to ±0.6 for visualisation). (D) Top 50 enriched Reactome pathways (when ranked by mean normalised enrichment score; NES) from gene set enrichment analysis of differentially expressed genes in Lyz2^+^ vs. Lyz2^−^ fibroblasts. Significantly enriched pathways (adjusted p<0.05) are coloured, with colour representing NES (trimmed to ±3 for visualisation). Volcano plots show differentially expressed genes from bulk RNA-sequencing of STZ vs. control mice bone marrow-derived macrophages, either (E) unstimulated or (F) stimulated with interferon-γ and lipopolysaccharide. Key fibroblast genes are highlighted.

To investigate whether bone marrow cells from STZ treated mice would also display fibroblast-like features, or a greater propensity to produce myeloid-derived fibroblasts, we examined previously generated bulk RNA-seq data from bone marrow-derived macrophages of STZ mice vs. control mice (Edgar et al., 2021). We found that unstimulated bone marrow-derived macrophages from STZ mice showed an upregulation of the fibroblast markers *Vim* and *Col4a2* compared to bone marrow-derived macrophages generated from control mice (Figure 4E). Furthermore, stimulation with interferon-γ and lipopolysaccharide resulted in increased expression of genes associated with myeloid cell to fibroblast transition: *Acta2*, *Tgfb1* and *Stat6* (Yan et al., 2016) specifically in bone marrow-derived macrophages from STZ treated mice (Figure 4F).

### Decorin is dysregulated across cells and is linked with human type 1 diabetes

Our analysis indicated the organ of origin, rather than disease group, as the primary driver of cell-type-specific gene expression profiles. To investigate functional features that are shared across cell-types and organs, we used gene set enrichment analysis and identified commonly dysregulated pathways, including ECM-related processes (e.g. ‘ECM proteoglycans’, ‘Collagen formation, ‘Extracellular matrix organization’), mRNA splicing/processing (e.g. ‘mRNA splicing’, ‘metabolism of RNA’), and heat shock function (e.g. ‘Cellular response to heat stress’) (Figure S7).

We also noted that a small number of genes were commonly, and highly differentially, regulated on organ-level analysis in STZ-treated mice. To identify these genes, we ranked all genes based on the frequency of significant differential expression across organs and cell types of STZ-treated mice (Figure 5A). We hypothesised that these genes might be relevant to the systemic effects of hyperglycaemia and in particular four genes were upregulated across many cell types and organs in STZ mice: the proteoglycan decorin (*Dcn)*, gelsolin (*Gsn*), matrix Gla protein (*Mgp*) and metallothionein 1 (*Mt1*). Gelsolin is a cytoskeletal regulator of actin filament assembly/disassembly, whilst decorin and matrix Gla protein are both components of the extracellular matrix. Fittingly, decorin (in addition to the proteoglycan biglycan; *Bgn*, and multiple collagen species) was among the leading-edge genes predicted to drive upregulation of the ‘ECM proteoglycans’ gene set across multiple cell types (Figure S7).

**Figure 5.**
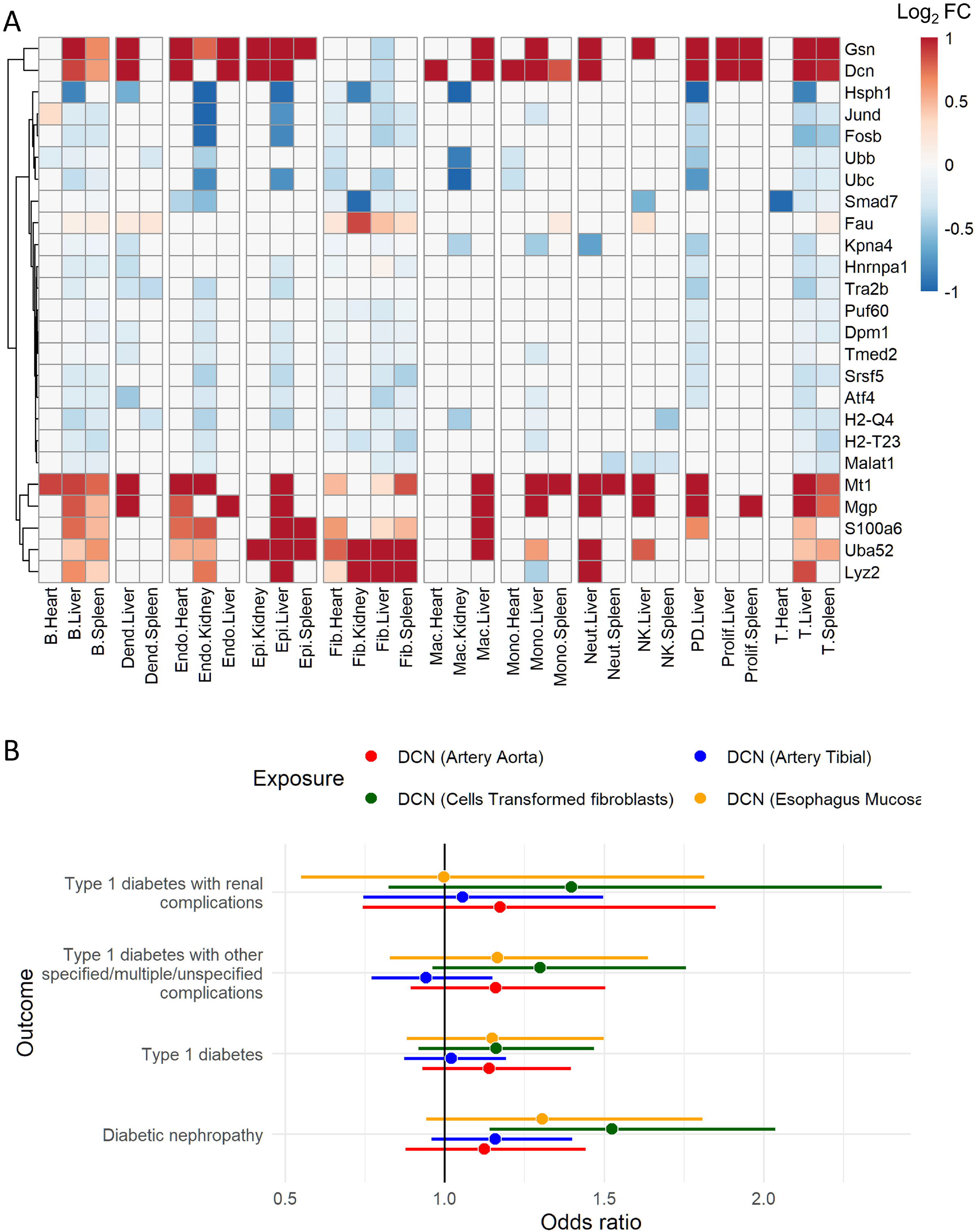
Decorin is dysregulated across cells and is linked with human type 1 diabetes. (A) The top 25 genes as ranked based on the frequency of significant differential expression (STZ vs. control) across all organ/cell types (for those with >150 total cells). Significantly differentially expressed genes (adjusted p<0.05) are coloured, with colour representing log2FC (trimmed to ±1 for visualisation). (B) Association of eQTLs for decorin with human type 1 diabetes traits was assessed by two-sample Mendelian randomisation. Odds ratio is plotted with bars representing lower and upper confidence intervals. DCN, decorin; Dend, myeloid dendritic cell; Endo, endothelial cell; Epi, epithelial cell; Fib, fibroblast; Mac, macrophage; Mono, monocyte; Neut, neutrophil.

Given the prominence of decorin, we sought to interrogate its association with complications in human diabetes. We performed a two-sample Mendelian randomisation analysis using *cis* eQTLs for decorin as instrumental variables and T1D / T1D with complications traits from the FinnGen GWAS study (Kurki et al., 2023) as outcomes (Figure 5B) and associated an eQTL regulating decorin transcription in fibroblasts with risk of diabetic nephropathy (Figure 5B).

### Cellular communication indicates fibroblast activation

STZ treatment resulted in perturbed gene expression across cell-types and organs. To understand how these changes affect the organ-specific interplay of immune and ‘non-immune’ cells, we assessed cellular crosstalk based on the expression of known ligands and receptors. We determined the number of interactions between selected cell types (Figure 6A) and the interaction strength (adjusted according to cell number; Figure 6B) using the STZ treatment data. To assess changes in cellular cross talk, we next compared the interaction strength between cell types of STZ-treated and control mice (Figure 6C). We detected altered cell communication from and to fibroblasts in each organ of STZ mice. Furthermore, STZ treatment resulted in increased signalling from ECs to fibroblasts, macrophages, T cells and ECs, likely driven by the ligand-receptor pair of Col4a2 and the plasma membrane proteoglycan, syndecan-4 (Sdc4; Figure S8). Our interaction analysis also revealed several high strength signalling relationships in the liver of STZ mice, particularly from fibroblasts to most other cell types. Overall, these data indicate the increased activation (via cellular communication) of fibroblasts upon induction of hyperglycaemia.

**Figure 6.**
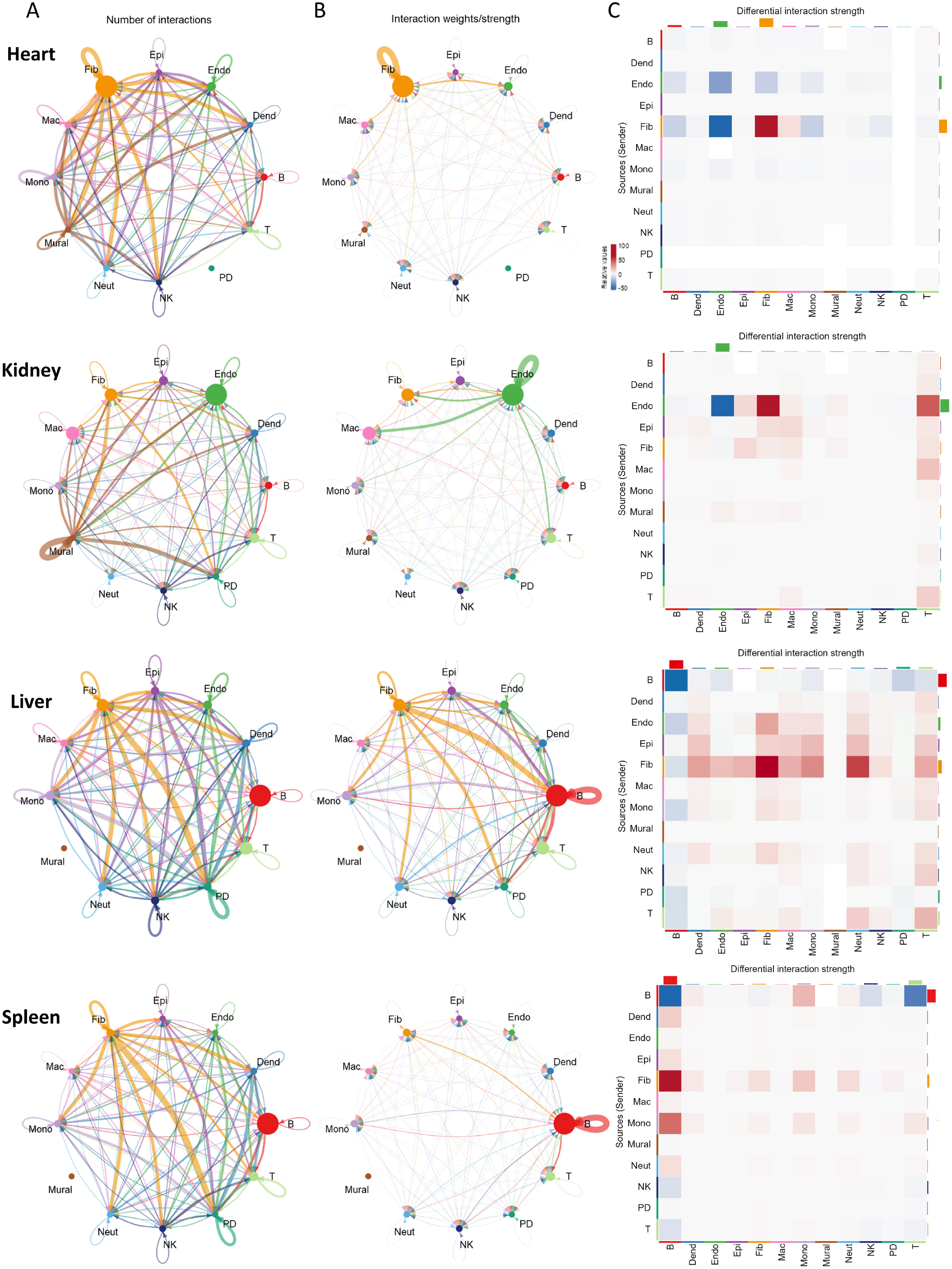
Cellular communication indicates fibroblast activation. Signalling interaction scores were evaluated by expression of ligand/receptor pairs in cells within each organ using CellChat and circle diagrams represent (A) interaction number and (B) interaction strength (adjusted according to cell number in the STZ condition). (C) The predicted strength of interactions between cell types per organ was then compared between STZ and control mice and summarised as a heatmap, with colours representing differential interaction strength. Dend, myeloid dendritic cell; Endo, endothelial cell; Epi, epithelial cell; Fib, fibroblast; Mac, macrophage; Mono, monocyte; Neut, neutrophil, PD; plasmacytoid dendritic cell.

## Discussion

Diabetes and hyperglycaemia lead to complications affecting multiple organs. The pathological processes underlying these complications are plausibly mediated by common cell types such as ECs, monocytes, macrophages and fibroblasts. We hypothesised that early responses in these cell types, before organ-specific pathologies manifest, might be shared, and we reasoned that this could be important since it would suggest targets for preventive and therapeutic interventions. Combining scRNA-seq profiling with a mouse model of type 1 diabetes and an early timepoint after hyperglycaemia induction, we identified dysregulation of pathways and cellular function of plausible relevance to the complications of hyperglycaemia. However, according to the single-cell transcriptome profiles, most cells were aligned according to organ of origin, rather than glycaemic status. Fibroblasts were the exception in this regard, and we identified the following disease-relevant features: (1) Across organs, fibroblasts from hyperglycaemic mice shared numerous functional characteristics, many of which were characteristics suggestive of an activated state. (2) We identified a high abundance subset of *Lyz2*^+^ fibroblasts, of likely myeloid origin and comprising almost half of the fibroblasts in the liver and kidney of STZ mice. Furthermore, (3) we found evidence of endo- and epithelial to mesenchymal transition in addition to markers of endothelial dysfunction. (4) Mediators of ECM remodelling, including decorin, were highly upregulated in many non-fibroblast cells, and Mendelian randomisation based on SNPs that determine decorin expression in fibroblasts showed a role in renal fibrosis. Finally, (5) a role for fibroblasts as early effectors in hyperglycaemia is supported by cell interaction modelling, in which inferred ligand-receptor pairings also suggest a central role for fibroblasts.

A switch to myofibroblast phenotype occurs in T2D (Fowlkes et al., 2013) and myofibroblasts have been linked to progression of diabetic nephropathy (Fowlkes et al., 2013; Pedagogos et al., 1997). We found upregulation of myofibroblast markers and p53-regulated metabolic pathways in STZ mouse fibroblasts, particularly in the heart and liver, in addition to an increased number of high-mitochondria fibroblasts in the STZ mouse heart. We also found that STZ mouse fibroblasts expressed high levels of lysozyme 2 (*Lyz2)*, a marker of myofibroblasts originating from a myeloid lineage (Guerrero-Juarez et al., 2019). These *Lyz2*^+^ fibroblasts exhibited increased expression of contractile markers (e.g. beta-actin) as well as monocyte (*Cd14*) and macrophage (*Cd68*) markers. Furthermore, *Lyz2^+^* fibroblasts had increased expression of genes associated with M1 polarised macrophages, including MHC class II genes and *Il1b*, which suggests that M1 macrophages could be a source of *Lyz2^+^* fibroblasts. To support this hypothesis, we analysed bulk transcriptomic data of bone marrow-derived macrophages from STZ mice (Edgar et al., 2021) and found that unstimulated STZ macrophages showed an upregulation of classical fibroblast markers (e.g. vimentin and *Col4a2*), and these genes were also expressed by a majority of *Lyz2*^+^ fibroblasts in the current study. Furthermore, stimulation of bone marrow-derived macrophages from STZ mice under M1-polarising conditions resulted in the upregulation of *Acta2*, *Tgfb1* and *Stat6* – genes that have been associated with transition of myeloid cells to fibroblasts (Yan et al., 2016). Fibroblast-like cells or fibrocytes originating from haematopoietic progenitors (rather than mesenchymal progenitors) have been detected in multiple contexts (Ogawa et al., 2006). For example, circulating monocytes can differentiate to fibrocytes and migrate to wound sites to assist in repair (Abe et al., 2001) and fibroblast-like cells derived from macrophages have also been implicated as an important driver of wound repair (Goren et al., 2009; Mirza et al., 2009; Sinha et al., 2018). In the context of diabetes, incomplete/reduced conversion of macrophages to fibroblasts can impair wound repair (Sinha et al., 2018). Conversely, dysregulation or over-activation of wound repair processes are also implicated in progressive fibrotic diabetic complications including diabetic nephropathy (Kanasaki et al., 2013). Furthermore, macrophage-derived myofibroblasts are drivers of collagen deposition in renal fibrosis (Meng et al., 2016) and cardiac fibrosis (Cieslik et al., 2011).

The increased number of infiltrating myeloid-origin myofibroblasts/fibrocytes and resulting ‘dilutional effect’ also likely led to the observed overall reduction in mRNA of collagen-coding genes and ECM mediators in STZ mouse fibroblasts. Any functional significance of an apparent reduction in collagen mRNA averaged across all fibroblast subtypes is not clear, as collagen genes remained highly expressed in the majority of STZ mouse fibroblasts. In any event there is likely to be a disconnect between collagen mRNA and protein levels, as post-transcriptional regulation may be more important than transcriptional regulation for type I collagen accumulation in the context of fibrosis (Stefanovic, 2013). There will be an inevitable disconnect between collagen mRNA and protein levels, as STZ mice develop histopathological signs of nephropathy, including fibrosis, at the later e.g. 6 month timepoints (Glastras et al., 2016; Sugimoto et al., 2007).

Interestingly, we found increased expression of collagen and ECM-related genes in other non-fibroblast cell types, including heart and liver ECs, which also had upregulated vimentin, a key marker of EndoMT/EMT. Transition of ECs to myofibroblasts has been shown to promote renal fibrosis in STZ mice (Li et al., 2009). Furthermore, lineage tracing experiments demonstrated that at 1 month post STZ treatment, EC-derived myofibroblasts are more numerous in kidney interstitium (around 10% of all myofibroblasts at 1 month, and 24% at 6 months after STZ) and this is associated with increased collagen type IV protein (Li et al., 2009). Myofibroblasts from ECs undergoing EndoMT have also been implicated as pro-fibrotic drivers of proliferative diabetic retinopathy (Abu El-Asrar et al., 2015), and EndoMT has been observed in diabetic heart and aorta ECs (Zhao et al., 2021). Interestingly, we found an increased expression of SCA1 (*Ly6a*) in heart fibroblasts, kidney ECs and liver epithelial cells isolated from our STZ-treated mice. SCA1 is linked to mesenchymal stemness/self-renewal/plasticity (Xin et al., 2005) and SCA1-expressing cardiac fibroblasts have been linked to heart failure (Chen et al., 2018). Together, these data further implicate the plasticity of myeloid cells, ECs and epithelial cells as sources of fibroblasts in response to early hyperglycaemia/T1D, and as a precursor to the later stage fibrosis associated with diabetic complications.

In addition to dysregulated fibroblast gene expression, we also found several genes whose expression were altered across different cell types upon STZ treatment. These included decorin, a small leucine-rich proteoglycan that is an inhibitor of TGFβ and an important regulator of fibrosis (Border et al., 1992; Hildebrand et al., 1994). Decorin expression was upregulated in cells from STZ mouse heart, kidney, liver and spleen. Previous work suggests that decorin might be a protective factor induced by diabetes. A variant allele of decorin was associated with slower progression of diabetic nephropathy in T1D patients (Cosmo et al., 2002). Further, studies using decorin knockout mice found increased nephropathy/proteinuria in the STZ model (Merline et al., 2009; Williams et al., 2007). However, using a Mendelian randomisation experiment, we associated a decorin eQTL in human fibroblasts with increased risk of diabetic nephropathy, suggesting that species- and cell type-specific effects should be considered when targeting decorin for therapeutic intervention.

We found several cellular changes in STZ mice that were reminiscent of pathogenic processes involved in diabetic complications, particularly in the kidney. This included altered expression of genes associated with EC dysfunction, such as hypoxia/heat shock and angiogenic factors, in STZ mouse ECs. We noted strong downregulation of heat shock protein-encoding genes (e.g. *Hsph1*, *Hspa1a, Hspb1*) and associated heat shock-related Reactome pathways (e.g. ‘Cellular response to heat stress’, ‘Regulation of HSF1-mediated heat shock’). Defective responses to hypoxia and/or cellular stress are implicated in hyperglycaemia-induced EC dysfunction (Fadini et al., 2019), a cellular state that can inhibit the induction of heat shock proteins which have been described as a protective against hyperglycaemia-induced cellular damage (Bellini et al., 2017). We also detected lower expression of plasminogen activator (*Plat*) and *Icam1* in ECs of STZ kidneys in accordance with previous work on EC dysfunction (Abebe and Mozaffari, 2010; Hadi and Suwaidi, 2007). Impaired angiogenesis is another aspect of EC dysfunction that underlies diabetic microvascular and macrovascular complications (Fadini et al., 2019) and fibroblasts are important promoters of angiogenesis via secretion of growth factors including VEGF (Newman et al., 2011). Notably, we found that STZ mouse fibroblasts had downregulated *Vegfa* expression in kidney and liver. Differential splicing regulation in kidney proximal tubule has also been linked to a pro-inflammatory phenotype in diabetes, implicating altered splicing in diabetic kidney disease (Wu et al., 2022). We found several pathways relating to RNA processing (‘mRNA splicing’, ‘Metabolism of RNA’, ‘Processing of Capped Intron-Containing Pre-mRNA’) and genes coding for spliceosome regulator serine/arginine-rich splicing factor 5, 6 and 9 (*Srsf5, Srsf6, Srsf9*) downregulated in multiple STZ mouse cells, including kidney ECs.

Previous work has shown that STZ-induced hyperglycaemia causes steatosis with raised hepatic triglyceride with infiltrating monocytes, neutrophils and Tregs in the liver (Lee et al., 2016). While we did not detect differences in leukocyte cell type abundance by scRNA-seq in the liver, we did observe increased liver mass and differential leukocyte transcription, including upregulation of the inflammatory complement genes *C1qa* and *C1qb* in STZ mouse liver monocytes/macrophages. Activation of the complement system is particularly important in metabolic tissues such as the liver (Phieler et al., 2013) and has been linked to pathogenesis of diabetes and its associated vascular complications (Shim et al., 2020), and we also noted upregulated complement genes *C1qa*, *C1qb* and *C1qc* in *Lyz2*^+^ fibroblasts of STZ mouse hearts. Experimental hyperglycaemia has been linked to enhanced myelopoiesis (Flynn et al., 2020; Nagareddy et al., 2014) and specifically within the STZ model (Nagareddy et al., 2013). We also found upregulation of the myelopoiesis-associated genes *S100a6*, *S100a8* and *S100a9* (Flynn et al., 2020; Nagareddy et al., 2014, 2013) in STZ mouse macrophages and monocytes (and notably, of *S100a8* in *Lyz2*^+^ spleen fibroblasts).

There were some limitations to this study. First, the number of mice per group affected statistical power and therefore the capacity to detect differences, e.g. in cell proportion between groups. Smaller groups also meant that greater sensitivity in detecting differential gene expression was obtained from individual cells rather than from pseudo-bulk profiles. Second, the cell types captured were likely to be affected differently by experimental methodology and thus the proportions measured were not necessarily a precise reflection of the cellular makeup of tissues *in vivo*. Third, STZ has been suggested to have direct hepatotoxic effects including apoptosis of hepatocytes (Kohl et al., 2013). We did not observe STZ-induced toxic effects on hepatocytes (annotated here as epithelial cells within the liver), but detected fewer B-cells in livers and spleens from STZ-treated mice, which may be associated with previously described direct B-cell toxicity of STZ (Muller et al., 2011).

Overall, we used an unbiased systemic approach to identify a number of cell-type-specific and organ-specific genes as well as functional pathways as potential drivers of diabetes-associated complications. Most notably, fibroblasts shared patterns of differential gene expression across organs and were either of monocyte / macrophage origin or derived by ‘reprogramming’ of other cell types via epi- and endo-mesenchymal transition. Interestingly, these processes were set in place early in response to hyperglycaemia and before overt organ pathology, suggesting that early intervention might be beneficial (or even necessary) to ameliorate some complications of diabetes. Therefore, our findings have translational relevance because fibrosis is a common feature of multiple organ pathologies that complicate type 2 diabetes. Based on our data, future studies will explore the mechanisms by which hyperglycaemia drives cell type transition providing potential targets for therapeutic intervention. Recent evidence from our group and others has shown that elevated blood glucose drives myelopoiesis (Edgar et al., 2021; Flynn et al., 2020) and induces trained immunity that had sustained effects on macrophage function and pathogenicity (Edgar et al., 2021). The current findings direct future interest towards the role of hyperglycaemia in driving fibroblast activation and functional programming, including directly or indirectly from the bone marrow. Finally, our study provides a rich resource for researchers to investigate the biological processes underlying early diabetic disease.

## Materials and methods

### Animals

All animal protocols were conducted in accordance with the UK Home Office under the guidance of the operation of the Animals (Scientific Procedures) Act 1986, complying to all ethical regulations and the institutional review board guidelines (project license 30/3374; January 27, 2016). Wild-type C57BL/6J (12–14 weeks old) male mice were randomly assigned to control or STZ experimental groups. Diabetes was induced by intraperitoneal injection of low dose (42–45 mg·kg−1·/d−1) STZ over 5 consecutive days. 8 weeks after the final streptozotocin injection, n=3 control mice and n=4 STZ mice with nonfasted blood glucose levels >20 mmol/L were euthanised, and tissue was collected.

### Cell isolation

Heart (ventricle), liver and kidneys were dissected and finely minced on ice using scissors. Cells were enzymatically dissociated in collagenase II (500U/ml; Thermo Fisher 17101015) in 10 ml HBSS, with agitation at 37°C for 1 hour. Dissociated heart, liver and kidney cells were passed through a 70 µm cell strainer and rinsed with 3 ml ice-cold HBSS. Spleens were passed through a 70 µm cell strainer into 1X Red Cell Lysis Buffer (1.5 mM ammonium chloride, 100 µM sodium bicarbonate, 11 µM EDTA disodium) and incubated for 10 minutes at room temperature. All cells were centrifuged for 5 min at 400 g at 4°C, then resuspended in 10% v/v DMSO in foetal bovine serum (FBS), aliquoted and frozen at −80°C before sequencing.

### 10X single cell RNA sequencing

Cells were revived by thawing at 37 °C for 1 min in a water bath and slowly transferred into 10 ml tubes filled with 9 ml pre-warmed RPMI containing 10% FBS and centrifuged at 500 g for 5 min. Next, cells were resuspended in 500 µl RPMI containing 10% FBS and labelled with commercially available DNA-labelled antibodies (TotalSeq-A, Biolegend) at 4 °C for 30 min. Following incubation, cell suspensions were washed three times with PBS containing 2 % BSA. After the final wash, cell suspensions were resuspended in PBS containing 2 % BSA and a viability dye (Zombie Red, Biolegend) for discrimination between live and dead cells. Next, 10,000 viable cells were sort-purified from the four organs of each individual mouse using a SH800 cell sorter (Sony). The four cell fractions were pooled for processing as a single sample according to the manufacturer’s protocol, while the antibody-linked barcodes enabled sample-specific demultiplexing of the sequencing data. scRNA-seq libraries were generated using the Chromium Controller and the Next GEM Single Cell 3’ Reagent Kit (v3, 10x Genomics) according to the manufacturer’s instructions. Libraries were sequenced by the Biomedical Sequencing Facility at the CeMM Research Center for Molecular Medicine of the Austrian Academy of Sciences, using the Illumina NovaSeq 6000 platform.

### Single cell library analysis

Reads from UMIs were aligned and counted per gene with 10X Genomics Cell Ranger 5.0.1 (Zheng et al., 2017) and the Seurat 4.1.0 R-package (Butler et al., 2018; Hao et al., 2021; Stuart et al., 2019) was used for HTO demultiplexing and doublet removal. Low quality and potential doublet cells were excluded by presence of two or more HTO tissue assignments, high percent mitochondria (>25%), number of genes detected (<300 or >5000) and number of UMIs detected (>25,000). Normalisation was performed for each sample with sctransform 0.3.3 (Hafemeister and Satija, 2019) using the Gamma-Poisson method provided by glmGamPoi 1.8.0 (Ahlmann-Eltze and Huber, 2020), with regression for percent mitochondria and predicted cell cycle difference (i.e. S-score minus G2M-score).

### Integrated single cell analysis

All single cell transcriptome profiles were integrated with Seurat using the ‘SCT’ method to minimise batch effects. Variation of gene expression across cells was assessed by PCA and then UMAP dimensional reduction based on the first 30 dimensions. Cell clusters were determined using Seurat with a resolution of 0.8, which was found to be optimal for cluster resolution and stability. Next, we used the Seurat ‘FindConservedMarkers’ function to identify marker genes of the individual clusters that were conserved across STZ and control groups. Cell types were annotated based on marker genes with a combined iterative approach including manual database searches of Tabula Muris (Schaum et al., 2018) and the Mouse Cell Atlas (Han et al., 2018), literature searches and the automated tools scCATCH 3.1 (Shao et al., 2020) and SingleR 1.10.0 (Aran et al., 2019). Pseudo-bulk RNA-seq profiles were generated by summing reads per gene across all cells per sample followed by calculating gene expression with DESeq2 1.36.0 (Love et al., 2014) and normalising across genes with a variance stabilising transformation.

### Inter-disease group single cell comparisons

Differential gene expression was assessed between STZ and control cells of each general cell type per tissue using the ‘FindMarkers’ function of Seurat with the Wilcoxon rank sum test, with significant adjusted p-values determined at p < 0.05. All genes from each comparison were ranked by sign(log2FC) x -log10(p-value) and evaluated using gene set enrichment analysis with fGSEA 1.22.0 (Korotkevich et al., 2021) to determine enrichment of mouse mapped versions of the Reactome pathway annotation set (version 0.3) from MSigDB (Subramanian et al., 2005), limited to pathways with < 200 genes. Enriched gene sets were collapsed to remove non-independent gene sets. Over-representation analysis was performed using the hypergeometric test within the fGSEA package for distinct sets of up-regulated and down-regulated differential genes (log2FC > 0.1, p < 0.05) per tissue/cell type (for those with > 150 cells) and for shared genes between cells of combinations of tissues. Cell signalling within each tissue was assessed by expression of ligand/receptor pairs using CellChat 1.0.0 (Jin et al., 2021) with adjustment for the effect of cell population size.

### Mendelian randomisation analysis

Two-sample Mendelian randomisation analyses were carried out using the TwoSampleMR R-package provided by MR-Base (Hemani et al., 2018, 2017). Instrumental variables were eQTL loci from the GTEx database (Aguet et al., 2017) and outcomes were T1D traits from the FinnGen GWAS study (Kurki et al., 2023).

### Statistical analyses

Statistical analyses and visualisations were performed with R 4.1.1. The statistical significance threshold for testing was set at p < 0.05 and appropriate adjustment for multiple testing (Benjamini Hochberg or Bonferroni) was utilised for all analyses.

## Sources of funding

This work was supported by research grants from the British Heart Foundation (BHF) Centre of Research Excellence, Oxford (RE/13/1/30181 and RE/18/3/34214); British Heart Foundation Project Grant (PG/18/53/33895) and a British Heart Foundation Intermediate Fellowship (FS/IBSRF/22/25110); the Tripartite Immunometabolism Consortium, Novo Nordisk Foundation (NNF15CC0018486 and NNF20SA0064144); Oxford Biomedical Research Centre (BRC).

## Supplementary figure legends

**Figure S1. QC and demultiplexing.** (A–C) Scatter plots of all cells with lines illustrating QC filters for percent mitochondrial reads (percent.mt), total number of reads (nCount_RNA) and number of unique genes detected (nFeature_RNA). Doublets and negatives were determined by presence of >1, or 0, hashtag oligo (HTO) tags, respectively. (D–G) Ridge plots represent log2 ‘expression level’ (i.e. number of HTO reads) per cell, summarised per demultiplexed organ. (H) t-distributed stochastic neighbour embedding (tSNE) plot of cells clustered by HTO expression, indicating singlet clusters and intermediate doublets (with two different HTO tags, e.g. HTO-Liver_HTO-Spleen).

**Figures S2-S5. Differentially expressed genes and pathways.** Volcano plots showing differences in gene expression in key cell types (fibroblasts, endothelial cells, monocytes and macrophages), for those with >150 total cells, in heart (Figure S2), liver (Figure S3), kidney (Figure S4) and spleen (Figure S5). Points were highlighted based on adjusted p-value (p < 0.05) and fold change (log2FC > 0.5). Bar plots indicate gene set enrichment analysis of Reactome pathways within ranked genes from each cell type. Significantly enriched pathways are shown with bars indicating normalised enrichment score.

**Figure S6. Relative cell abundance.** Plots indicate relative frequency of cell sub-clusters per organ, split by disease group. Frequency of STZ vs. control mice cells was assessed by t-test. *, p<0.05; ***, p<0.001. Dend, myeloid dendritic cell; Endo, endothelial cell; Epi, epithelial cell; Fib, fibroblast; Mac, macrophage; Mono, monocyte; Neut, neutrophil, PD; plasmacytoid dendritic cell.

**Figure S7. Key enriched pathways across cell types and organs.** Key enriched Reactome pathways from gene set enrichment analysis across organs/cell types are highlighted, representing common functions for extracellular matrix (ECM), mRNA processing and heat shock. Significantly enriched pathways (adjusted p<0.05) are coloured, with colour representing normalised enrichment score (trimmed to ±2 for visualisation). Endo, endothelial cell; Fib, fibroblast; Mac, macrophage; Mono, monocyte.

**Figure S8. Upregulated ligand-receptor pairs in hyperglycaemia.** Ligand-receptor pairs were assessed by CellChat within differentially expressed genes in STZ vs. control mouse cells. Highlighted key ligand-receptor pairs are predicted to drive increased signalling from STZ mouse kidney endothelial cells. Sector width within the chord diagram represents total predicted signalling strength for an interaction. Dend, myeloid dendritic cell; Endo, endothelial cell; Epi, epithelial cell; Fib, fibroblast; Mac, macrophage; Mono, monocyte; Neut, neutrophil, PD; plasmacytoid dendritic cell.

## Notes

### Competing Interest Statement

The authors have declared no competing interest.

