## Supplementary methods for "Multi-organ single-cell RNA-sequencing reveals early hyperglycaemia responses that converge on fibroblast dysregulation"

### Supplementary data

#### 1 Identification of cell clusters and sub-clusters

Six distinct B cell clusters were identified based on expression of the markers *Cd79a* and *Cd79b* (B CD21, B Mt.Hi, B naïve 1, B naïve 2, B plasma and B spleen). One cluster expressed *Cr2* (CD21; B CD21) and two clusters had particularly high expression of *Ccr7* and *Cd83*, a marker of naïve B cells (Lüthje et al., 2008) (B naïve 1 and B naïve 2). A single cluster of plasma B cells was identified based on expression of the marker *Jchain* (Castro and Flajnik, 2014) and lower expression of *Cd79a* and *Cd79b* than in other B cell clusters (B plasma). A single B cell cluster was annotated based on high proportion (mean >10%) of mitochondrial UMIs (B Mt.Hi). Another B cell cluster was specifically identified within spleen and was designated as such (B spleen). Two clusters of dendritic cells were identified as expressing the dendritic cell marker *Itgax*. One of these clusters highly expressed *S100a4* (labelled Dend s100a4) whereas the other cluster had low expression of *S100a4* (Dend). Three highly distinct clusters were determined to be ECs from heart (Endo H), liver (Endo L) and kidney (Endo K). All expressed the EC marker *Pecam1* (Figure 1C). Cells from clusters Endo H and Endo K had high expression of *Ly6a*, while Endo L cells did not. Three clusters of epithelial cells were identified using the marker *Car2* (Epi 1, Epi 2, Epi 3). One of these specifically expressed *Car3* (Epi 2), and two expressed the markers *Cdh1* and *Epcam* (Epi 1 and Epi 3). Six fibroblast clusters specifically expressed the markers *Col1a2* and *Mmp2* (Fib, Fib act, Fib aWNT, Fib Il6, Fib Mt.Hi and Fib SCA1). One of these fibroblast clusters had particularly higher expression of *Il6* (Fib Il6). Another had relatively high expression of the mesenchymal stemness/plasticity marker *Ly6a* (Challen et al., 2009) (SCA1; Fib SCA1). Two fibroblast clusters specifically expressed the fibroblast activation marker *Postn* (Muhl et al., 2020) (Fib act and Fib aWNT) and one of these also expressed *Wif1*, a marker associated with a subset of fibroblasts that suppresses WNT signalling (Muhl et al., 2020) (Fib aWNT). One of the fibroblast clusters was determined to have a higher mitochondria content based on high percentage (mean >10%) of mitochondrial UMIs and also had some, but lower expression of *Col1a2* and *Mmp2* than other fibroblast clusters (Fib Mt.Hi). Single macrophage and Kupffer cell clusters were found to express key marker *Cd68* and *C1qa* (Mac, Kupf). Macrophages were identified based on expression of the myeloid marker *Cd68* and complement component *C1qa,* with Kupffer cells distinguished based on their specificity to liver and expression of the resident macrophage marker *Vsig4* (Li et al., 2017), while non-resident macrophages did not express *Vsig4* but specifically expressed *Ms4a7*. Three monocyte clusters expressed *Cd68* in addition to *S100a4* (Mono, Mono Ly6 1, Mono Ly6 2). Two of these clusters expressed *Ly6c*, *Ccr2* and *Chil3* (Mono Ly6 1, Mono Ly6 2). One of the *Ly6c*+ monocyte clusters was differentiated from the other based on higher expression of *Cd14* and *Il1b* (Mono Ly6 2). A mural cell cluster was identified using the markers *Acta2* (α-SMA), *Des* and *Rgs5* (Smyth et al., 2018). However, due to low number of cells in this cluster, it was not possible to determine whether these were pericytes, vascular smooth muscle cells or other mural-like cells (or a combination of these cells). A single neutrophil cluster (Neut) was determined as expressing the markers *S100a8* and *S100a9*. A single natural killer cell cluster (NK) was identified using the markers *Nkg7* (Ng et al., 2020) and *Klrb1c* (Abel et al., 2018). One cluster of plasmacytoid dendritic cells were identified as expressing the marker *Ccr9* (Wendland et al., 2007). Four T cell clusters specifically expressed the marker *Cd3d*. Of these, two distinct clusters expressed *Cd8a* (T CD8 1, and to a lesser extent, T CD8 2) and *Nkg7*, two expressed *Cd4* (T CD4, T naïve) and two expressed *Ccr7* (T naïve, and to a lesser extent, T CD8 1).
