## Supplementary figures 1 for "Multi-organ single-cell RNA-sequencing reveals early hyperglycaemia responses that converge on fibroblast dysregulation"

### Figure S1

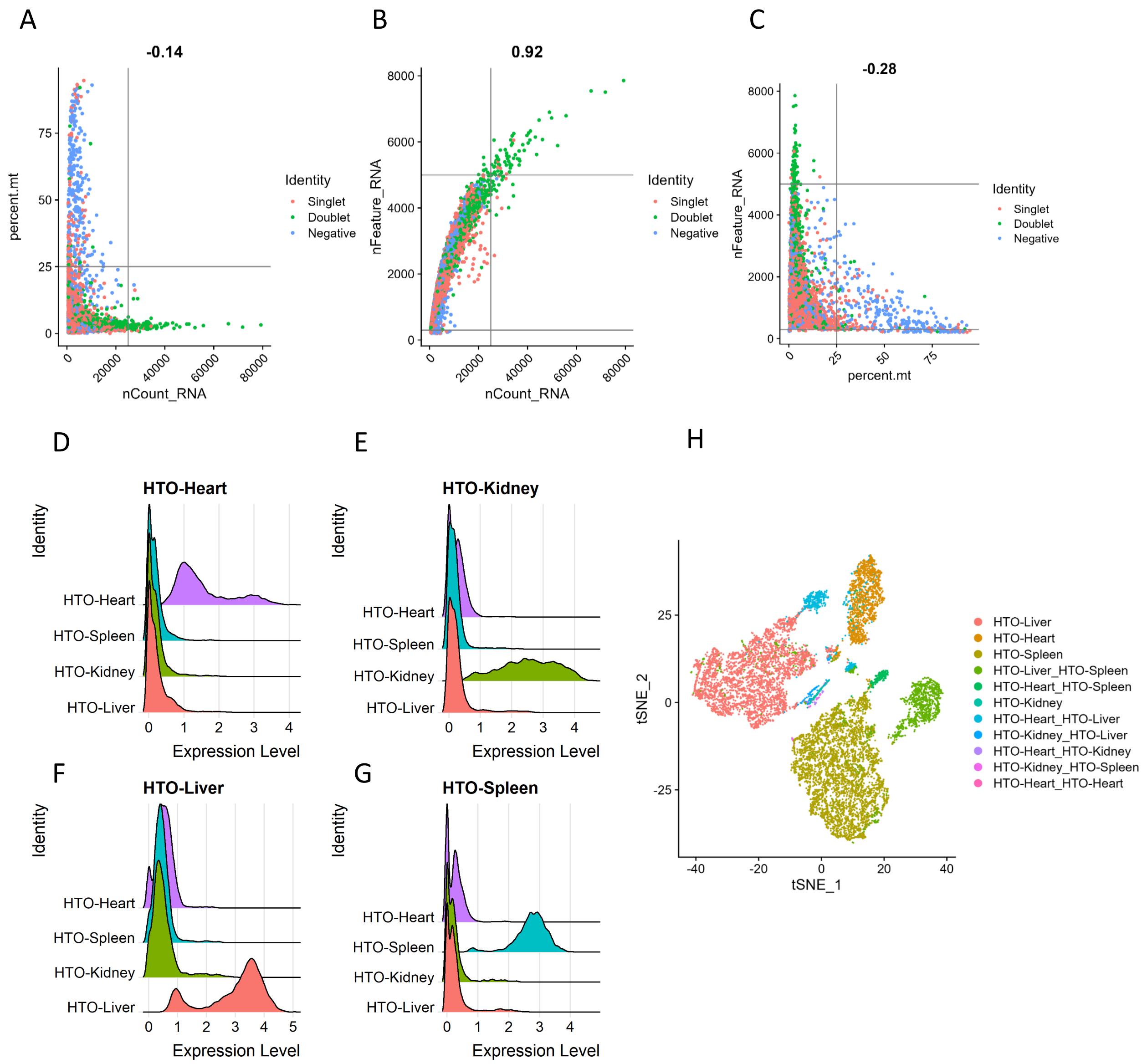
A
-0.14
percent.mt
nCount_RNA
Identity
Singlet
Doublet
Negative
B
0.92
nFeature_RNA
nCount_RNA
C
-0.28
nFeature_RNA
percent.mt
D
HTO-Heart
E
HTO-Kidney
F
HTO-Liver
G
HTO-Spleen
Identity
HTO-Heart
HTO-Spleen
HTO-Kidney
HTO-Liver
Expression Level
H
tSNE_2
tSNE_1
HTO-Liver
HTO-Heart
HTO-Spleen
HTO-Liver_HTO-Spleen
HTO-Heart_HTO-Spleen
HTO-Kidney
HTO-Heart_HTO-Liver
HTO-Kidney_HTO-Liver
HTO-Heart_HTO-Kidney
HTO-Kidney_HTO-Spleen
HTO-Heart_HTO-Heart

### Figure S2

#### Heart

##### Fibroblasts

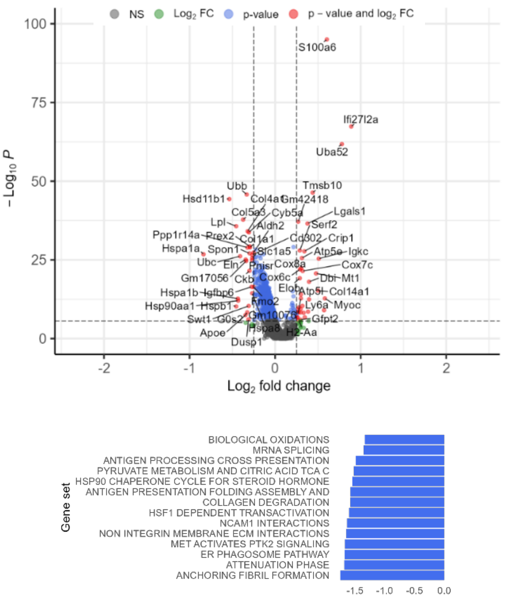


##### Endothelial cells

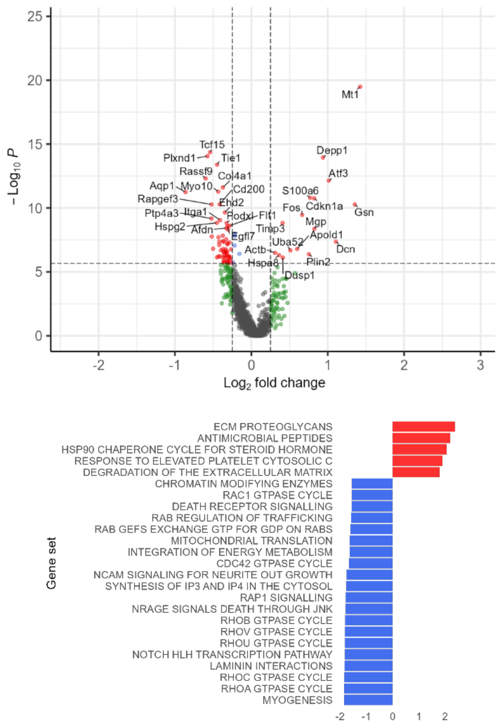


##### Monocytes

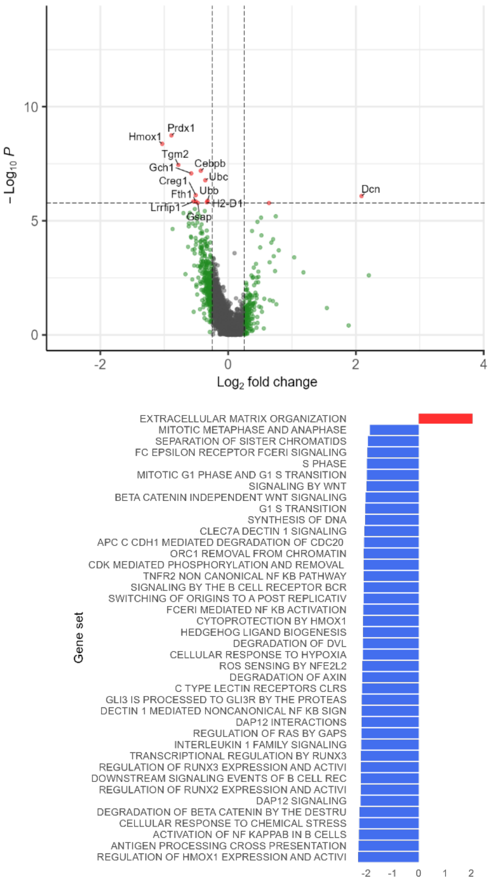


##### Macrophages

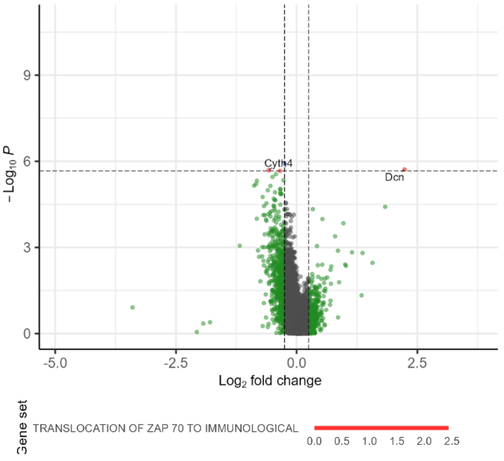


Pathway up in STZ
Pathway up in control

### Figure S3

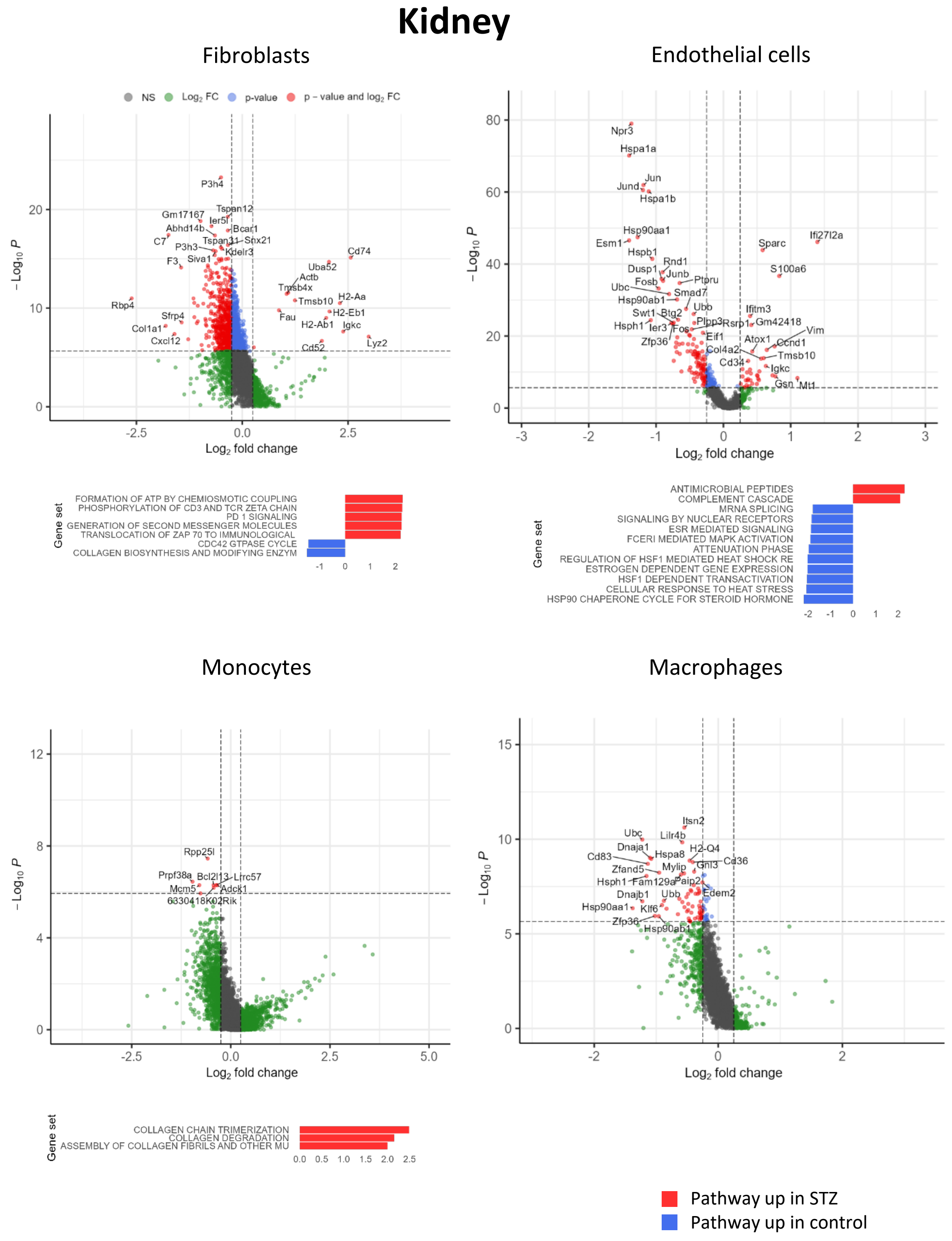
Kidney
Fibroblasts
Endothelial cells
NS
Log2 FC
p-value
p – value and log2 FC
Log2 fold change
– Log10 P
Gene set
FORMATION OF ATP BY CHEMIOSMOTIC COUPLING
PHOSPHORYLATION OF CD3 AND TCR ZETA CHAIN
PD 1 SIGNALING
GENERATION OF SECOND MESSENGER MOLECULES
TRANSLOCATION OF ZAP 70 TO IMMUNOLOGICAL
CDC42 GTPASE CYCLE
COLLAGEN BIOSYNTHESIS AND MODIFYING ENZYM
ANTIMICROBIAL PEPTIDES
COMPLEMENT CASCADE
MRNA SPLICING
SIGNALING BY NUCLEAR RECEPTORS
ESR MEDIATED SIGNALING
FCERI MEDIATED MAPK ACTIVATION
ATTENUATION PHASE
REGULATION OF HSF1 MEDIATED HEAT SHOCK RE
ESTROGEN DEPENDENT GENE EXPRESSION
HSF1 DEPENDENT TRANSACTIVATION
CELLULAR RESPONSE TO HEAT STRESS
HSP90 CHAPERONE CYCLE FOR STEROID HORMONE
Monocytes
Macrophages
COLLAGEN CHAIN TRIMERIZATION
COLLAGEN DEGRADATION
ASSEMBLY OF COLLAGEN FIBRILS AND OTHER MU
Pathway up in STZ
Pathway up in control

### Figure S4

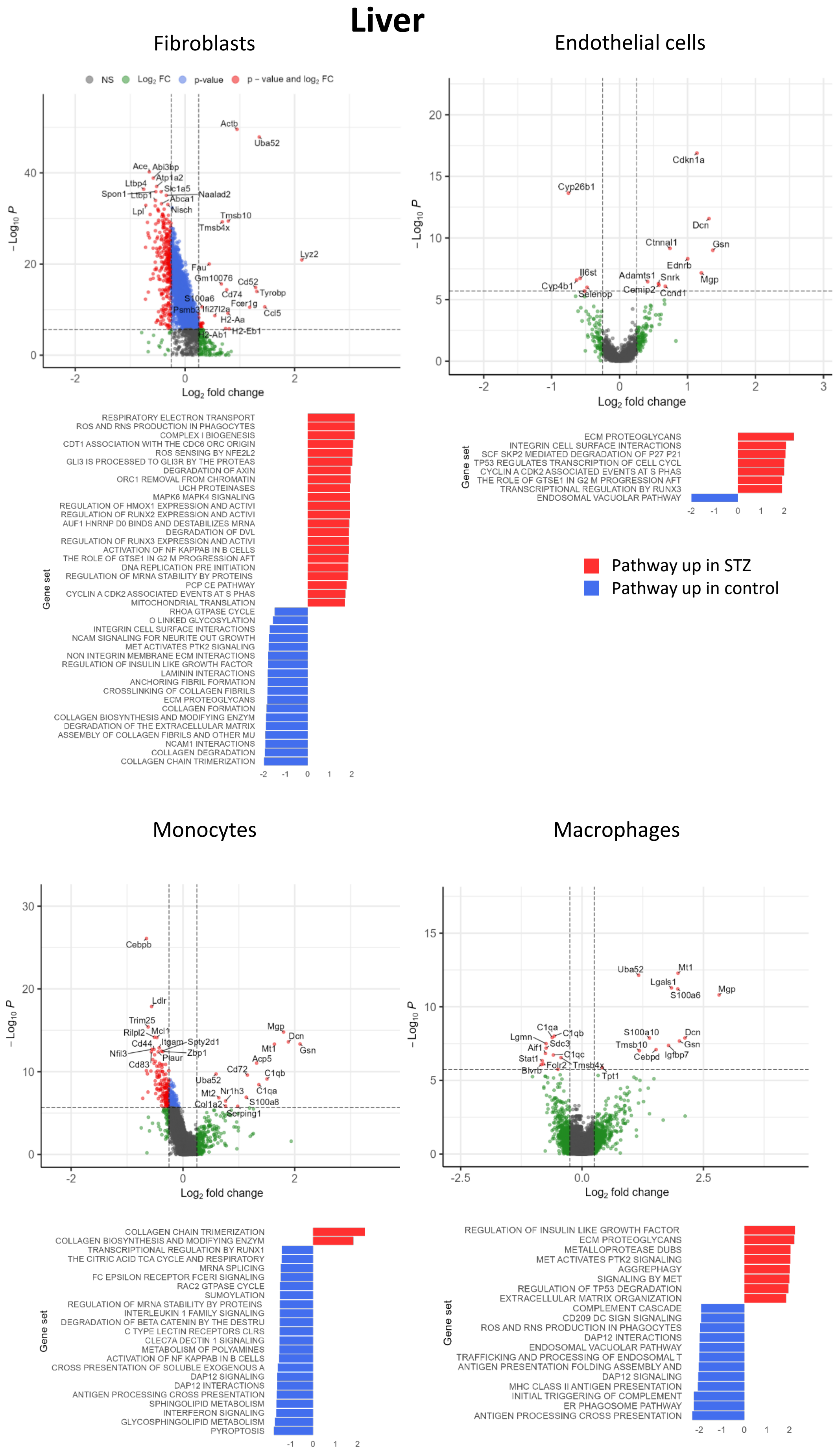

Liver
Fibroblasts
NS
Log2 FC
p-value
p – value and log2 FC
Actb
Uba52
Ace
Abi3bp
Atp1a2
Ltbp4
Slc1a5
Spon1
Ltbp1
Naalad2
Abca1
Nisch
Lpl
Tmsb10
Tmsb4x
Lyz2
Fau
Gm10076
Cd52
Cd74
Tyrobp
S100a6
Fcer1g
Psmb3
Ifi27l2a
Ccl5
H2-Aa
H2-Eb1
H2-Ab1
– Log10 P
Log2 fold change
RESPIRATORY ELECTRON TRANSPORT
ROS AND RNS PRODUCTION IN PHAGOCYTES
COMPLEX I BIOGENESIS
CDT1 ASSOCIATION WITH THE CDC6 ORC ORIGIN
ROS SENSING BY NFE2L2
GLI3 IS PROCESSED TO GLI3R BY THE PROTEAS
DEGRADATION OF AXIN
ORC1 REMOVAL FROM CHROMATIN
UCH PROTEINASES
MAPK6 MAPK4 SIGNALING
REGULATION OF HMOX1 EXPRESSION AND ACTIVI
REGULATION OF RUNX2 EXPRESSION AND ACTIVI
AUF1 HNRNP D0 BINDS AND DESTABILIZES MRNA
DEGRADATION OF DVL
REGULATION OF RUNX3 EXPRESSION AND ACTIVI
ACTIVATION OF NF KAPPAB IN B CELLS
THE ROLE OF GTSE1 IN G2 M PROGRESSION AFT
DNA REPLICATION PRE INITIATION
REGULATION OF MRNA STABILITY BY PROTEINS
PCP CE PATHWAY
CYCLIN A CDK2 ASSOCIATED EVENTS AT S PHAS
MITOCHONDRIAL TRANSLATION
RHOA GTPASE CYCLE
O LINKED GLYCOSYLATION
INTEGRIN CELL SURFACE INTERACTIONS
NCAM SIGNALING FOR NEURITE OUT GROWTH
MET ACTIVATES PTK2 SIGNALING
NON INTEGRIN MEMBRANE ECM INTERACTIONS
REGULATION OF INSULIN LIKE GROWTH FACTOR
LAMININ INTERACTIONS
ANCHORING FIBRIL FORMATION
CROSSLINKING OF COLLAGEN FIBRILS
ECM PROTEOGLYCANS
COLLAGEN FORMATION
COLLAGEN BIOSYNTHESIS AND MODIFYING ENZYM
DEGRADATION OF THE EXTRACELLULAR MATRIX
ASSEMBLY OF COLLAGEN FIBRILS AND OTHER MU
NCAM1 INTERACTIONS
COLLAGEN DEGRADATION
COLLAGEN CHAIN TRIMERIZATION
Gene set
Endothelial cells
Cdkn1a
Cyp26b1
Dcn
Ctnnal1
Gsn
Ednrb
Il6st
Adamts1
Snrk
Mgp
Cyp4b1
Cemip2
Selenop
Ccnd1
ECM PROTEOGLYCANS
INTEGRIN CELL SURFACE INTERACTIONS
SCF SKP2 MEDIATED DEGRADATION OF P27 P21
TP53 REGULATES TRANSCRIPTION OF CELL CYCL
CYCLIN A CDK2 ASSOCIATED EVENTS AT S PHAS
THE ROLE OF GTSE1 IN G2 M PROGRESSION AFT
TRANSCRIPTIONAL REGULATION BY RUNX3
ENDOSOMAL VACUOLAR PATHWAY
Pathway up in STZ
Pathway up in control
Monocytes
Cebpb
Ldlr
Trim25
Rilpl2
Mcl1
Cd44
Itgam
Spty2d1
Nfil3
Zbp1
Plaur
Cd83
Mgp
Dcn
Gsn
Mt1
Acp5
Cd72
C1qb
Uba52
Mt2
Nr1h3
C1qa
Col1a2
S100a8
Serping1
COLLAGEN CHAIN TRIMERIZATION
COLLAGEN BIOSYNTHESIS AND MODIFYING ENZYM
TRANSCRIPTIONAL REGULATION BY RUNX1
THE CITRIC ACID TCA CYCLE AND RESPIRATORY
MRNA SPLICING
FC EPSILON RECEPTOR FCERI SIGNALING
RAC2 GTPASE CYCLE
SUMOYLATION
REGULATION OF MRNA STABILITY BY PROTEINS
INTERLEUKIN 1 FAMILY SIGNALING
DEGRADATION OF BETA CATENIN BY THE DESTRU
C TYPE LECTIN RECEPTORS CLRS
CLEC7A DECTIN 1 SIGNALING
METABOLISM OF POLYAMINES
ACTIVATION OF NF KAPPAB IN B CELLS
CROSS PRESENTATION OF SOLUBLE EXOGENOUS A
DAP12 SIGNALING
DAP12 INTERACTIONS
ANTIGEN PROCESSING CROSS PRESENTATION
SPHINGOLIPID METABOLISM
INTERFERON SIGNALING
GLYCOSPHINGOLIPID METABOLISM
PYROPTOSIS
Macrophages
Uba52
Mt1
Lgals1
Mgp
S100a6
C1qa
Lgmn
C1qb
Aif1
Sdc3
S100a10
Dcn
Tmsb10
Gsn
Stat1
C1qc
Cebpd
Igfbp7
Folr2
Tmsb4x
Blvrb
Tpt1
REGULATION OF INSULIN LIKE GROWTH FACTOR
ECM PROTEOGLYCANS
METALLOPROTEASE DUBS
MET ACTIVATES PTK2 SIGNALING
AGGREPHAGY
SIGNALING BY MET
REGULATION OF TP53 DEGRADATION
EXTRACELLULAR MATRIX ORGANIZATION
COMPLEMENT CASCADE
CD209 DC SIGN SIGNALING
ROS AND RNS PRODUCTION IN PHAGOCYTES
DAP12 INTERACTIONS
ENDOSOMAL VACUOLAR PATHWAY
TRAFFICKING AND PROCESSING OF ENDOSOMAL T
ANTIGEN PRESENTATION FOLDING ASSEMBLY AND
DAP12 SIGNALING
MHC CLASS II ANTIGEN PRESENTATION
INITIAL TRIGGERING OF COMPLEMENT
ER PHAGOSOME PATHWAY
ANTIGEN PROCESSING CROSS PRESENTATION

### Figure S5

#### Spleen

##### Fibroblasts

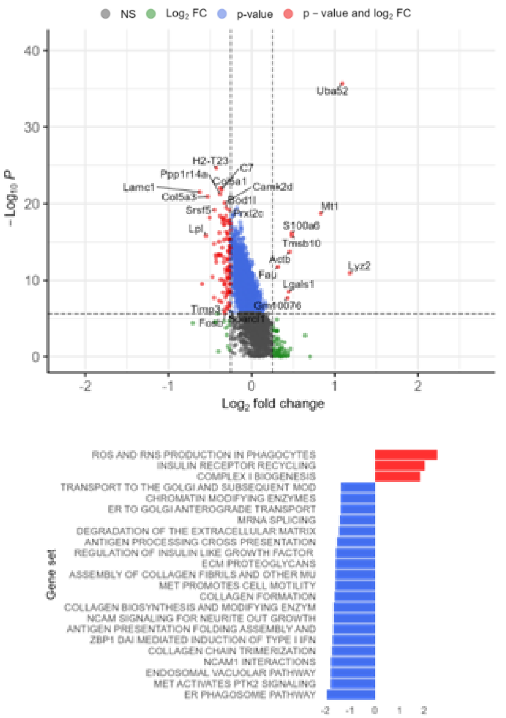


##### Monocytes

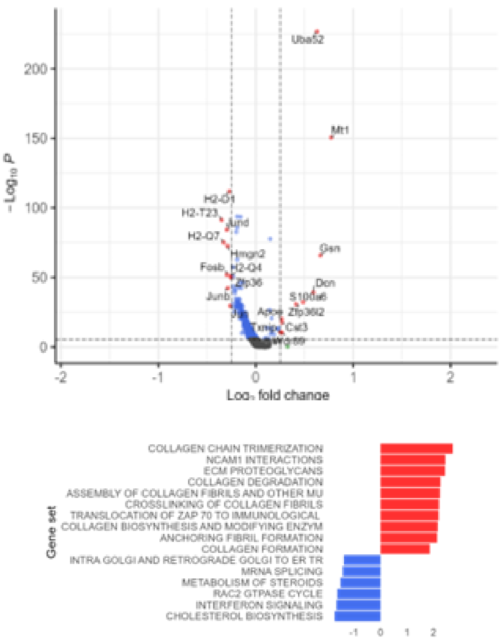


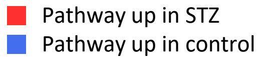


##### Macrophages

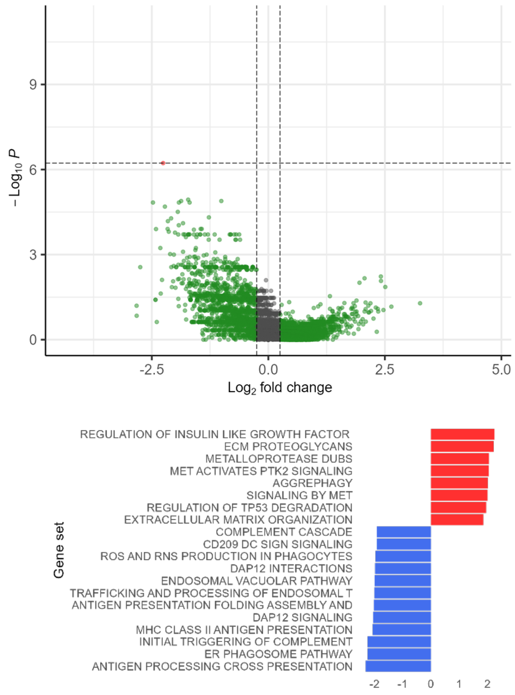

### Figure S7

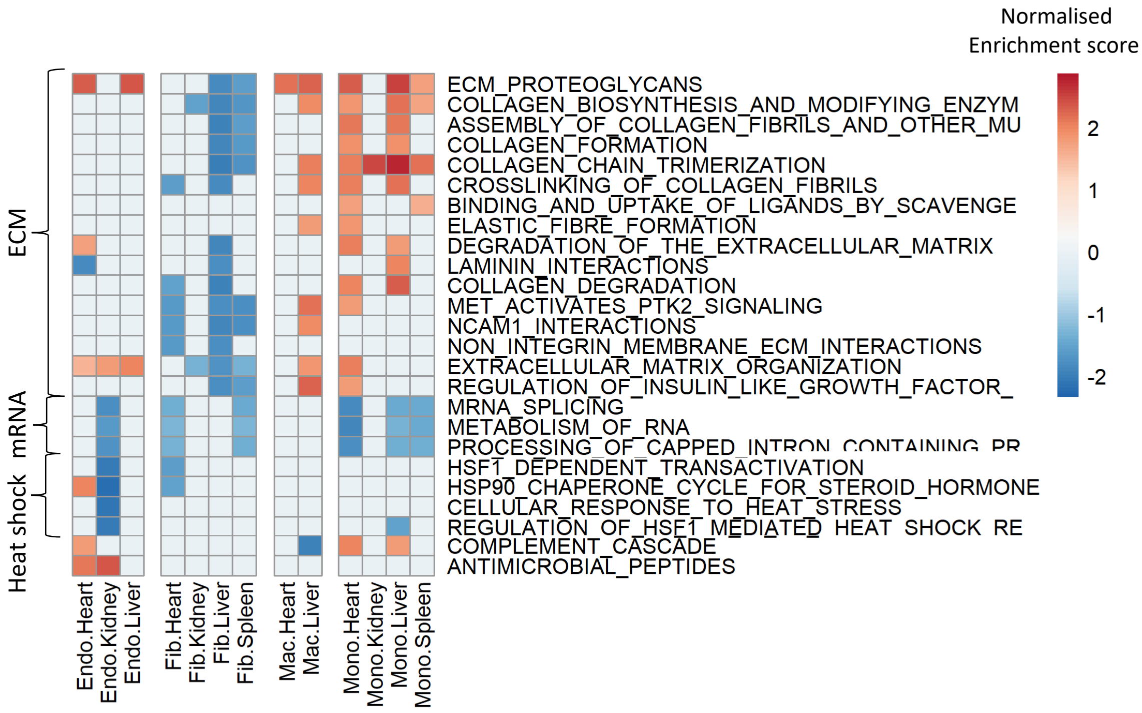
Normalised
Enrichment score
2
1
0
-1
-2
ECM
mRNA
Heat shock
ECM_PROTEOGLYCANS
COLLAGEN_BIOSYNTHESIS_AND_MODIFYING_ENZYM
ASSEMBLY_OF_COLLAGEN_FIBRILS_AND_OTHER_MU
COLLAGEN_FORMATION
COLLAGEN_CHAIN_TRIMERIZATION
CROSSLINKING_OF_COLLAGEN_FIBRILS
BINDING_AND_UPTAKE_OF_LIGANDS_BY_SCAVENGE
ELASTIC_FIBRE_FORMATION
DEGRADATION_OF_THE_EXTRACELLULAR_MATRIX
LAMININ_INTERACTIONS
COLLAGEN_DEGRADATION
MET_ACTIVATES_PTK2_SIGNALING
NCAM1_INTERACTIONS
NON_INTEGRIN_MEMBRANE_ECM_INTERACTIONS
EXTRACELLULAR_MATRIX_ORGANIZATION
REGULATION_OF_INSULIN_LIKE_GROWTH_FACTOR_
MRNA_SPLICING
METABOLISM_OF_RNA
PROCESSING_OF_CAPPED_INTRON_CONTAINING_PR
HSF1_DEPENDENT_TRANSACTIVATION
HSP90_CHAPERONE_CYCLE_FOR_STEROID_HORMONE
CELLULAR_RESPONSE_TO_HEAT_STRESS
REGULATION_OF_HSF1_MEDIATED_HEAT_SHOCK_RE
COMPLEMENT_CASCADE
ANTIMICROBIAL_PEPTIDES
Endo.Heart
Endo.Kidney
Endo.Liver
Fib.Heart
Fib.Kidney
Fib.Liver
Fib.Spleen
Mac.Heart
Mac.Liver
Mono.Heart
Mono.Kidney
Mono.Liver
Mono.Spleen

### Figure S8

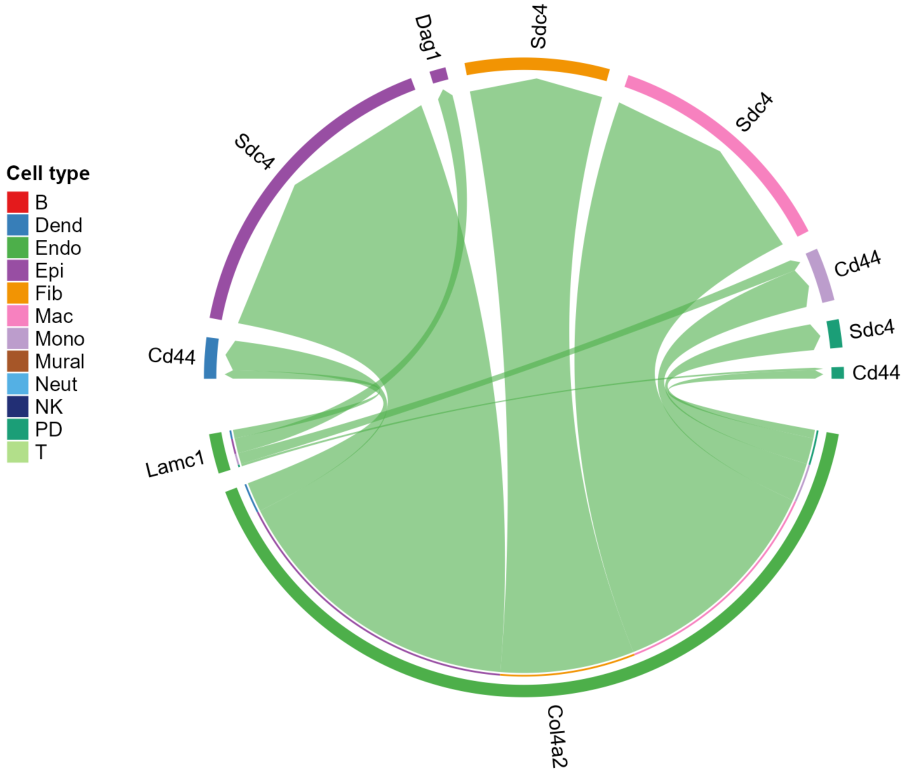
Cell type
B
Dend
Endo
Epi
Fib
Mac
Mono
Mural
Neut
NK
PD
T
Dag1
Sdc4
Sdc4
Sdc4
Cd44
Cd44
Sdc4
Cd44
Lamc1
Col4a2
