## Supplementary figures and images for "Multi-organ single-cell RNA-sequencing reveals early hyperglycaemia responses that converge on fibroblast dysregulation"

### Supplementary figures 2

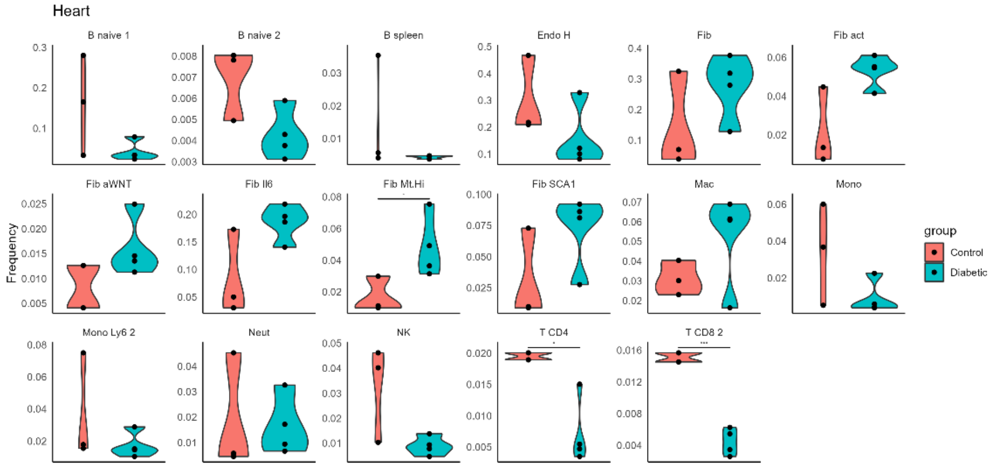

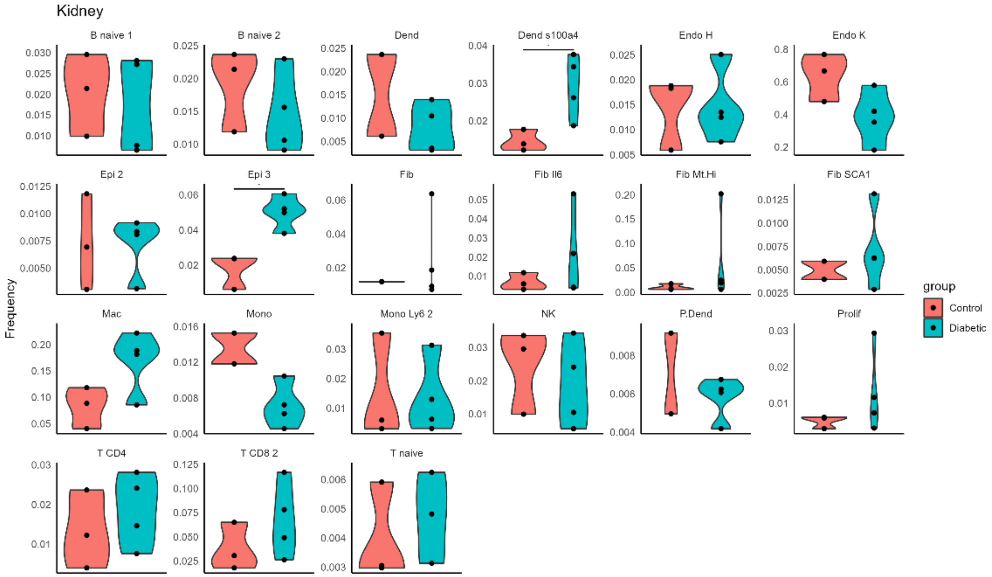

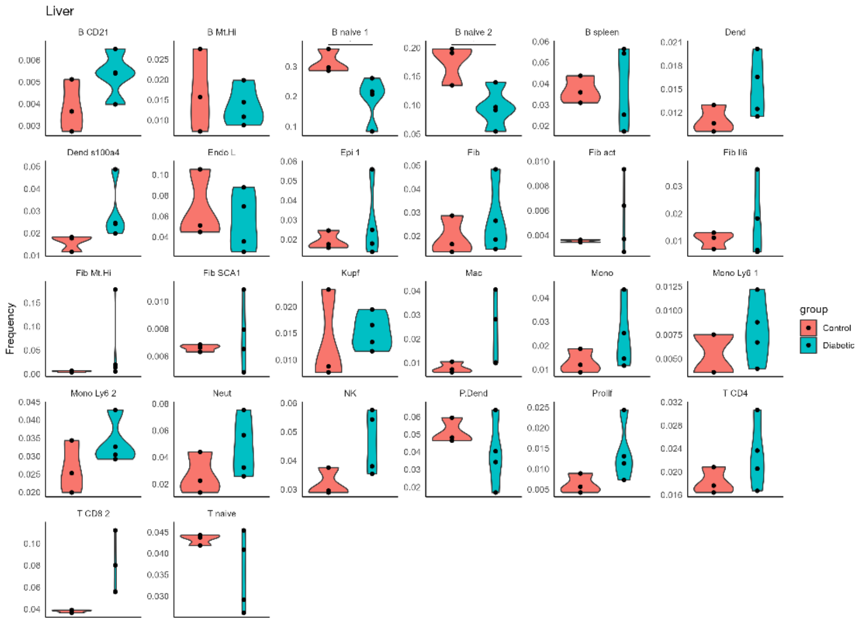

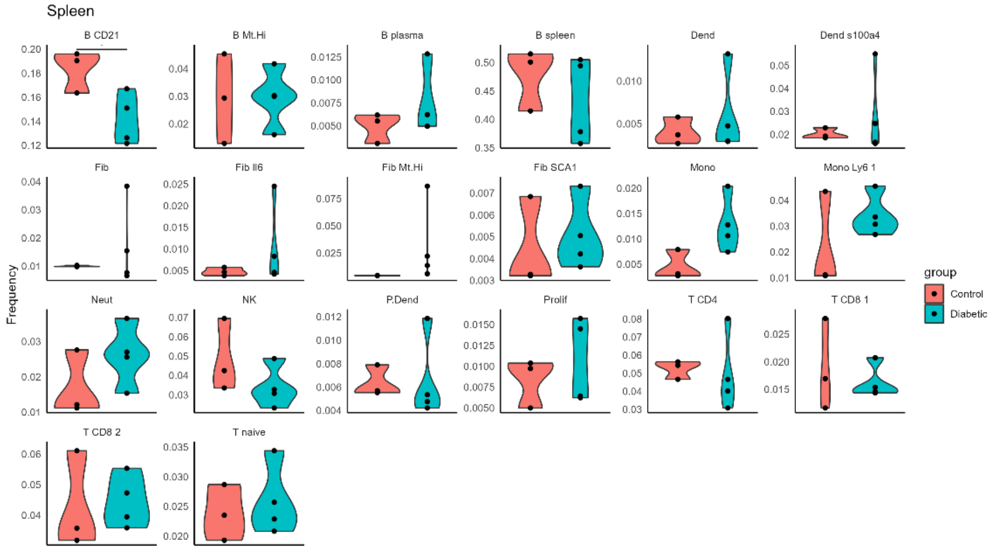
